# Unraveling the role of laminin(*lama5*) in maintenance of epithelial identity and polarity in bilayer zebrafish epidermis during development

**DOI:** 10.1101/2025.09.09.675236

**Authors:** Tooba Khan

## Abstract

Polarity is one of the fundamental properties of cells. It is characterised by the asymmetric distribution of lipids and proteins in the cell membrane and cortex, and organelles in the cell. Cell polarisation drives essential processes such as morphogenesis, cell migration, asymmetric cell division, directional transport of molecules across epithelium, nerve impulse transmission etc. Although, the importance of polarity proteins such as atypical protein kinase C (aPKC) and Lgl2 (lethal giant larvae 2) has been relatively well understood in the developing zebrafish epidermis, whether basal lamina components are important for polarity maintenance has remained unclear. I analysed the role of Lamininα5 (basal lamina component) in the establishment and maintenance of apicobasal polarity, in developing bilayered zebrafish epidermis. I found that the loss of laminin α5 function results in reduced E cadherin localization and increased cell spreading along with dynamic cell boundaries and increased cell proliferation indicating acquisition of mesenchymal traits. A similar phenotype was observed in integrin α6b mutant, which did not exacerbate in the double mutant embryos indicating that laminin α5 and integrin α6b function in the same pathway. Interestingly, periderm, the cell layer above basal epidermis, maintains its apicobasal polarity and epithelial integrity presumably via reinforcing the localization of aPKC and Lgl. My work unravels the importance of Laminin α5 and Integrin α6b interaction in the maintenance of epithelial characteristics in the basal layer of the developing zebrafish epidermis.

## Introduction

Polarisation of a cell refers to an asymmetric distribution of organelles, proteins, lipids, and cellular projections. It is one of the distinct features of epithelial cells, which are polarized along the apical-basal axis. They exhibit actin-based projections in the apical domain and polarized organization of junctions such as tight junctions, adherens junction, and desmosomes on the lateral side with tight junctions demarcating the apical domain and segregating it from the basolateral domain. The basal domain of the epithelial cells is in contact with the basal lamina formed of extracellular matrix proteins such as laminins, via integrin-based adhesion. The apical-basal polarity is controlled by four major complexes or modules viz. Par3-Par6-aPKC complex, Crumbs complex, Dlg-Scrib-Lgl module and Yurt-Cora-Na+K+ ATPase module. While Par and Crumbs complexes localised to the apical domain regulate the establishment and maintenance of the apical domain, Dlg-Scrib-Lgl and Yurt-Cora function as the regulators of the basolateral domain (Buckley & Jhonston, 2022). Establishment of apicobasal polarity is thought to occur spontaneously by self-recruitment of Par3/Bazooka to the apical membrane via its N-terminal oligomerization (CR1) domain, followed by recruitment and binding to the aPKC-Par6-Cdc42 as well as by stabilisation of apical Crumbs complex presumably via aPKC trans phosphorylation. This eventually leads to the formation of Par and Crumbs complexes in the apical domain of epithelial cells (Lin et al, 2000; Thompson, 2013). The apical and basolateral complexes act in a mutually antagonistic fashion to maintain their respective domains and hence the polarity in the epithelial cells (Chalmers et al., 2005, Tepass, 2012). Lipids exhibit asymmetric distribution along the apical basal axis of the epithelial cells too. For example, it has been shown that Phosphatidylinositol-4,5-bis phosphate (PtdIns(4,5)P2) is enriched in the apical domain, while Phosphatidylinositol-3,4,5-tris phosphate (PtdIns(3,4,5)P3) localization is augmented towards the basolateral side of the epithelial cell (Devergne et al., 2014; Gassama-Diagne et al., 2006; Martin-Belmonte et al., 2007; Raman et al., 2018). Importantly, PtdIns(4,5)P2 aids in recruitment of Annexin to the apical domain which eventually recruits aPKC and Par6 via CDC42 (Martin-Belmonte et al., 2007) suggesting that these phospho-inositides act upstream of polarity regulators.

Basement membrane is an integral part of the epithelial tissue and is formed of extracellular matrix proteins such as laminin, collagen, perlecan, nidogen, etc. (Matlin et al., 2017). Amongst these, laminins have been shown to play important role in regulating apicobasal polarity in 3D spheroid, and cell cultures of mammalian epithelial cells like MDCK (Akhtar and Streuli, 2013; Cohen et al., 2011; O’ Brien et al, 2001). Laminins are glycoproteins that are synthesized and secreted by epithelial cells towards the basal side. They are composed of three subunits-alpha (α), beta (β), and gamma (γ) that form heterotrimer in endoplasmic reticulum (ER) followed by post translational modification in Golgi, after which they are transported by microtubule towards the basal side for secretion (Bradshaw, 2016; Zajac & Horne-Bodanivic, 2022). Engagement of Integrins with Collagen matrix is linked to the activation of a Rho GTPase Rac1, basal actin organization and Laminin secretion (Lee and Streuli, 2014). A dominant negative form of Rac1 (N17Rac1, inactive GDP bound form of Rac1) leads to misassembled laminin and inversion of apicobasal polarity (O’ Brien et al., 2001). The polarized deposition and assembly of Laminin-1 (Li et al, 2002; Miner and Yurchenco, 2004) has been shown to be of the utmost importance for the formation of basement membrane (Yurchenco, 2004), to prevent an inversion of apicobasal cell polarity (Akhtar and Streuli, 2013; O’ Brien et al., 2001), and formation of limbs and neural tube (Miner et al., 1998). Furthermore, in MDCK 3D spheroid cultures, it has been shown that knockout of integrin β1 subunit, which functions as a laminin receptor, or its downstream effector ILK (Integrin linked kinases), lead to failure of polarity establishment and maintenance (Akhtar and Streuli, 2013; Yu et al., 2004) presumably via outside-in signaling. On the other hand, heterodimers Integrin α3β1, Integrin α6β1, Integrin α6β4 bind to laminins (Nishiuchi et al, 2006), and are required for laminin assembly (Matlin et al., 2017). Thus, the bidirectional signaling between the epithelial cells and the ECM regulates the organization of the basal lamina and the establishment and maintenance of the apico-basal polarity.

Whether and how laminins regulate the establishment and maintenance of the polarity in multilayered epithelia has remained unclear. Previous work has shown that Laminins begin to be localised towards the basal side of the developing zebrafish epidermis by 20 hours post-fertilisation. Furthermore, mutation in laminin α5 results in defects in the formation of pectoral fins and fin fold (Carney et al.,2010; Nagendran et al., 2015; Webb et al., 2007). However, its role in polarity regulation and its potential interactions with Integrins remain unaddressed. I found that in the absence of laminin α5 function, the basal cells show reduced E-cadherin localization, cell-cell detachment, dynamic cell protrusions, perturbed cell polarity (membrane Lgl2 levels decrease), and flatter cells (cell height decreases) suggesting loss of epithelial and acquisition of mesenchymal traits. Furthermore, the loss of Integrin α6b and simultaneous loss of laminin α5 and Integrin α6 results in a similar phenotype suggesting that both these genes function in the same pathway.

## Results

### 1. Laminin is highly expressed only in basal epidermis of zebrafish during development

Laminins are heterotrimers of glycoproteins that are deposited in the basement membrane (Matlin et. al., 2017). Laminin C1 or γ-1 is a component of Laminin-1 and other trimeric laminin complexes widely expressed in almost all basement membranes (Li et al., 2002; Matlin et al., 2017; Miner & Yurchenco, 2004; Yurchenco, 2011). Both 2D cell culture and 3D organoids cell culture studies of cell polarity and laminin have only a single cell layer (Akhtar and Streuli, 2013; Guo et al.,2024; O’ Brien et al., 2001; Yu et al., 2005). So far there are no studies that have shown laminin expression in an organised bilayered tissue context. Since zebrafish epidermis is composed of two layers during development, i.e., upper periderm layer and lower basal epidermis layer, I wanted to see which of these two or both layers express Laminin C1. To address this, I used Tg(lamc1:lamc1-sfGFPp1), a transgenic line that expresses the super folderGFP variant tagged to Laminin C1, under lamc1 promoter. I found that basal epidermis show high expression of Laminin C1-sfGFP at both 36hpf, and 48hpf (Fig 1a, 1b). Interestingly, periderm does not show Laminin C1-sfGFP expression at both stages (Fig S1). The region below basal epidermis is defined as the basement membrane region enriched in laminin isoforms α, β, γ which form the nascent or primary basement membrane sheet over which other basement membrane molecules incorporate to form a mature basement membrane structure (Hamill et al., 2009, Li et al., 2002; Li et al., 2003b; Matlin et. al., 2017). This region below basal epidermis, was found to be enriched in Laminin C1-sfGFP as well at both 48hpf, and 36hpf stages (Fig 1a. 1b). Thus, in a two-layered zebrafish epidermis, only basal epidermis in direct contact with the basement membrane express Laminin C1-sfGFP.

**Fig. 1.**
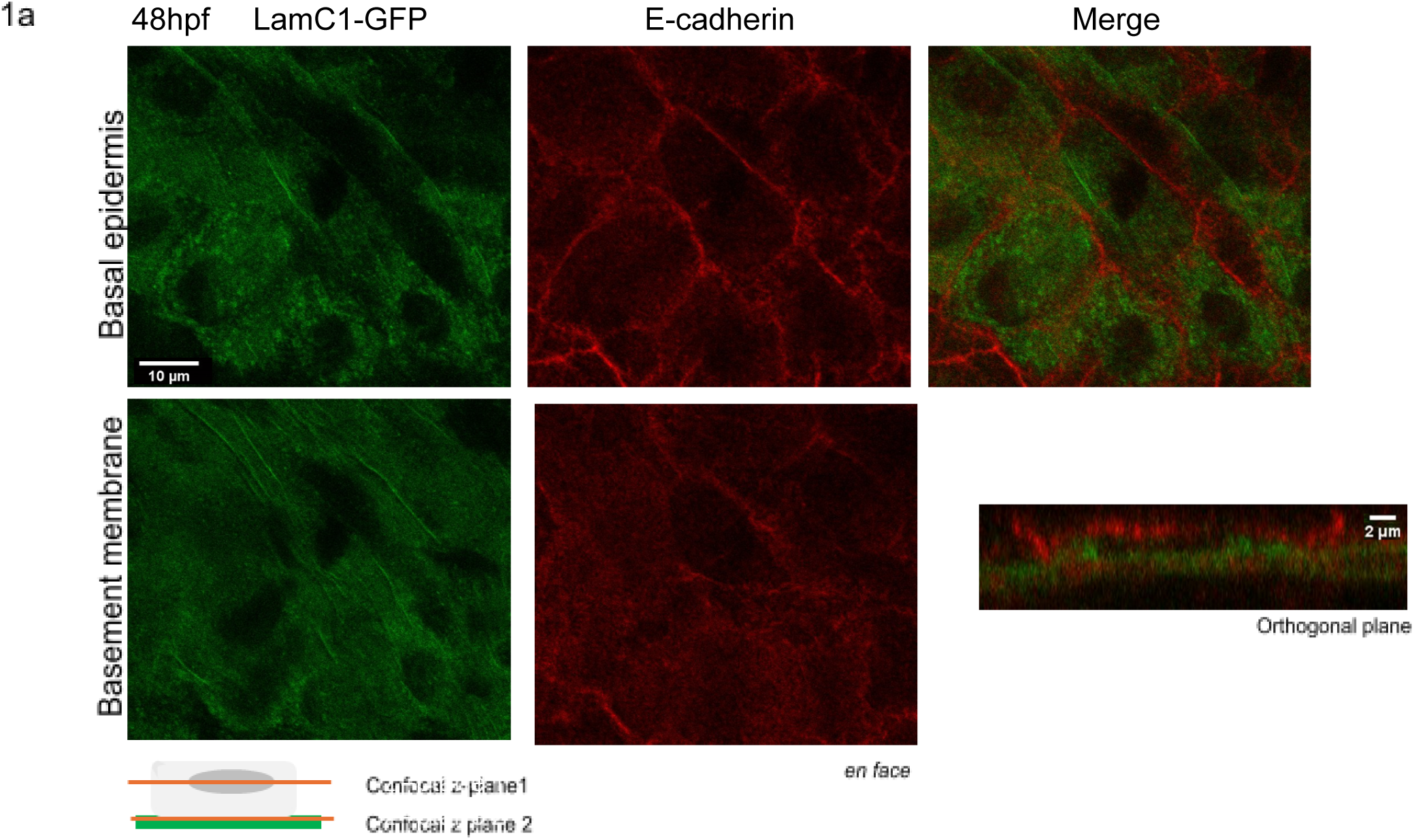
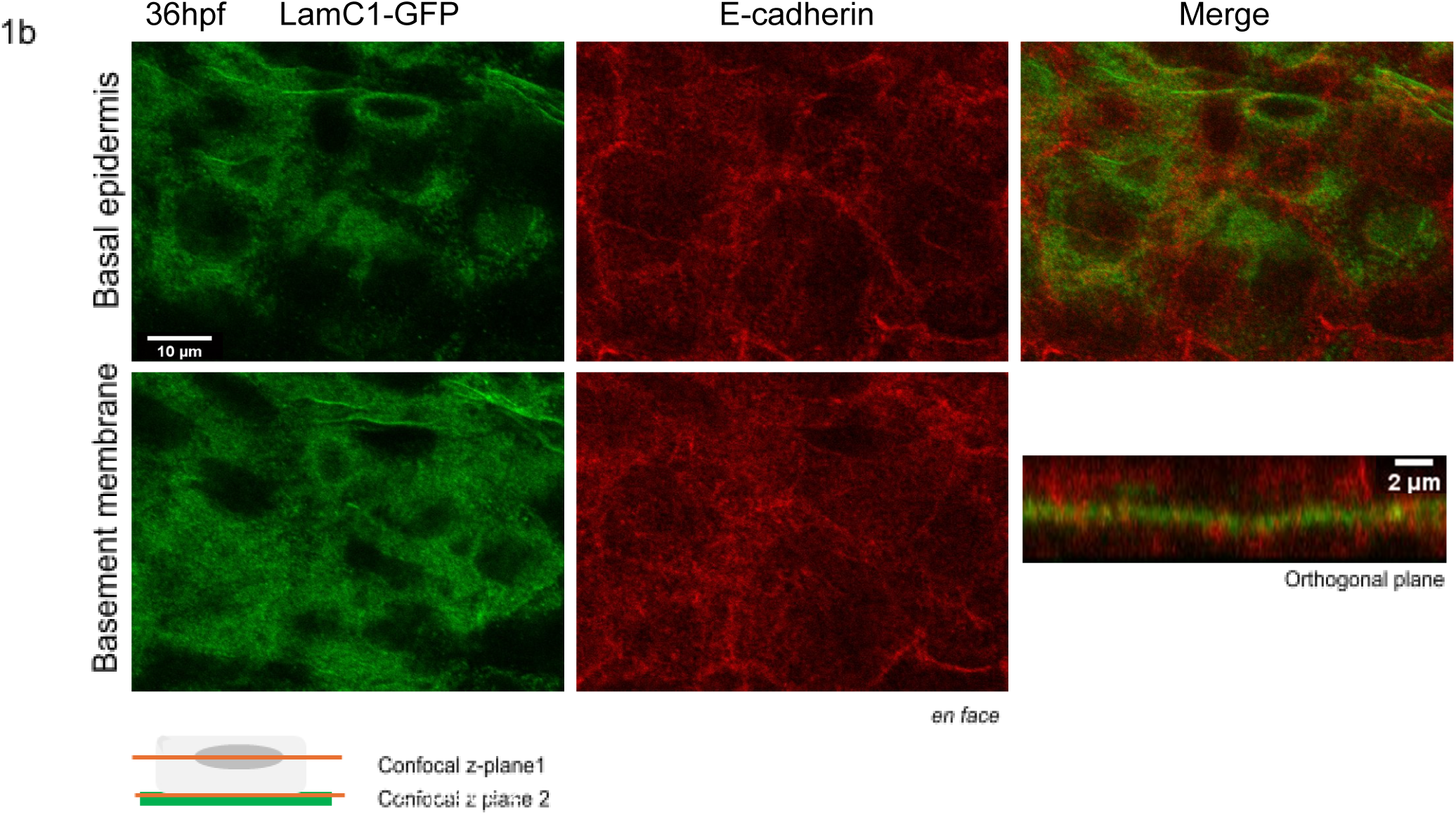
Basal epidermis expresses and is enriched in laminin gamma1 (*lamc1*). LamininC1-GFP was found to be expressed in basal epidermal cells marked by E-cadherin at 48hpf(Fig1a), and 36hpf(Fig1b). The region immediately below it (basement membrane region, lacking E-cadherin) was also enriched in LamininC1-GFP. The schematic diagram depicts one basal epidermal cell resting on a basement membrane towards its basal side. Confocal z-plane 1 lies inside the cell. Confocal z plane 2 lies just beneath when the cells disappear. (See supplementary Fig. S1). Scale bar depicts 10um length in *en face* images; 2um length in orthogonal plane. In images, LamC1-GFP is shown in green LUT, and membrane Lgl2 in red LUT (in Fiji).

### 2. Laminin α5 is required to maintain cell morphology and cell-cell adhesion protein E-cadherin levels at cell-cell junctions in basal epidermis

Despite laminin’s polarised localisation only towards the basal side of cells (Cohen et al., 2011, Roignot et al., 2013) there is very little understanding of its interplay with other cell polarity and junction proteins (Akhtar and Streuli, 2013; Daley et al., 2012; Guo et al.,2024; O’ Brien et al, 2001). I sought to understand if loss of laminin had any effect on cell polarity and junctions. Among other Laminin alpha isoforms, Laminin α5 was found to be highly enriched in zebrafish epidermis during early development (Danio cell dataset; Farrell et al., 2018; Sur et al., 2023). I used lama5 mutants (Webb et al., 2007; Zhang et al., 2022) to understand the effect of loss of Laminins in the basement membrane on cell junctions and apicobasal cell polarity. I found that loss of Laminin α5 in basement membrane results in reduced E-cadherin level at cell-cell junctions (Fig 2b, n=3, ****p<0.0001, Mann Whitney test) in comparison to their wild type siblings (WT sib). This might have an effect on cell-cell adhesion in basal epidermal cells. I then went on to see if loss of Laminin α5 had any effect on cell morphology of basal epidermis. I found a significant decrease in cell height of basal epidermis in lama5 mutants in comparison to their wild type siblings (Fig 2c, n=3, ****p<0.0001, Mann Whitney test). However apical perimeter of cells was largely unaffected in lama5 mutants (Fig 2d, n=3, ‘ns’p=0.5398, unpaired t test). Thus, loss of Laminin α5 causes reduction in cell-cell junction molecules E-cadherin levels that might have implications on cell-cell adhesion of basal epidermal cells, and also result in reduced cell height.

**Fig. 2.**
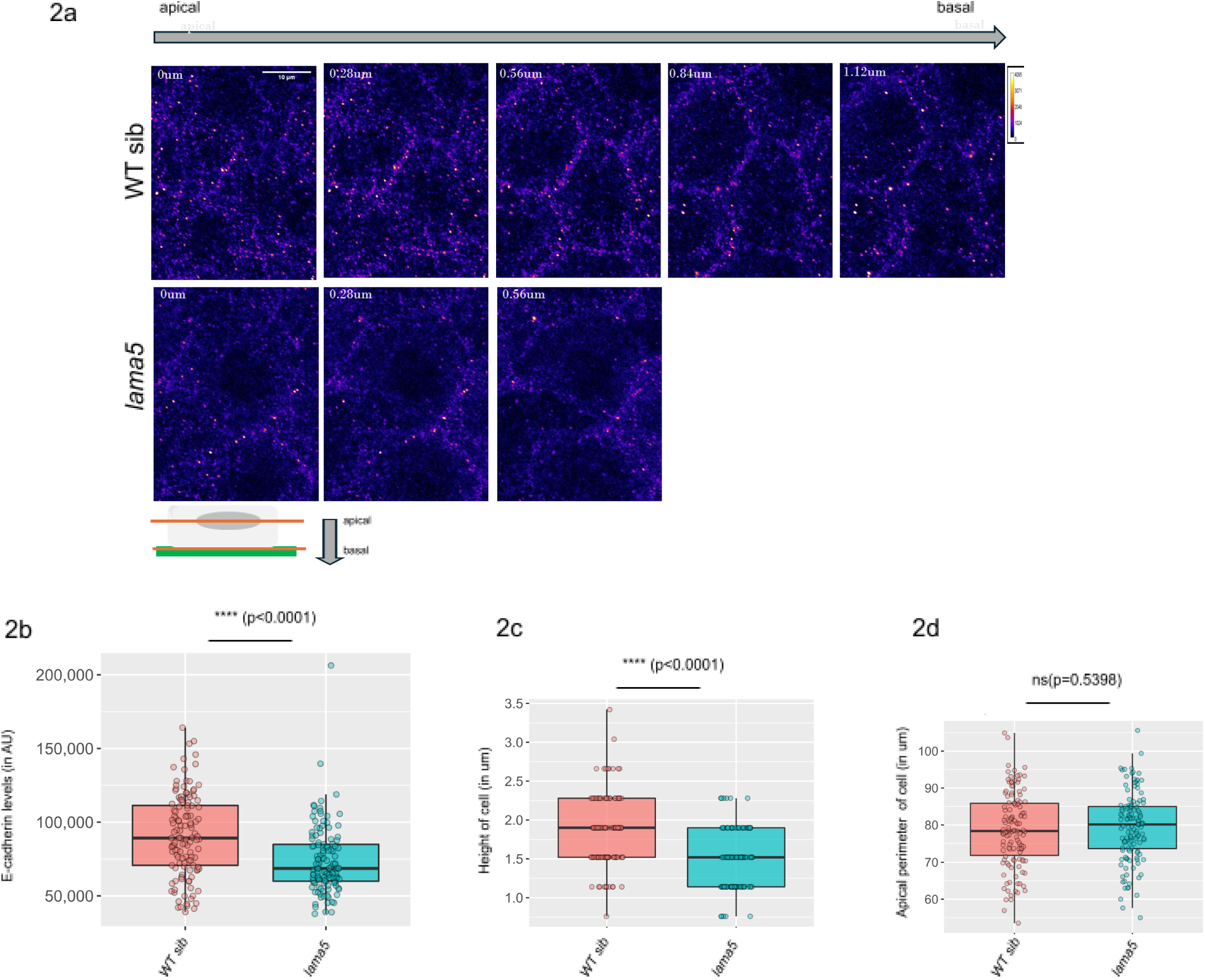
Junctional E-cadherin levels and cell morphology is affected in basal epidermis in *lama5* mutants. Total junctional E-cadherin levels goes down in basal epidermis upon loss of laminin alpha5 (Fig2a, 2b). Cell height decreases in basal epidermis in *lama5* mutants (Fig2c). Apical perimeter does not change significantly in *lama5* mutants (Fig2d). Fig 2b, 2c ****p<0.0001 (n=3, Mann Whitney test); Fig 2d ‘ns’p=0.5398 means (non significant difference, n=3, unpaired t test).The schematic diagram depicts one basal epidermal cell resting on a basement membrane towards its basal side. ‘apical’ for basal epidermis lies above close to periderm. ‘basal’ for basal epidermis lies below towards the basement membrane region. (See supplementary S2, and S14 for source file and statistical comparisons). Scale bar depicts 10um length. Images are shown as heat map with calibration bar ranging from 0 to 4095 (in Fiji).

### 3. Lgl2 basolateral membrane polarity protein localisation in basal epidermis depends on Laminin α5

Previous studies in zebrafish epidermis have shown the reciprocal effect of membrane Lgl2, and junctional E-cadherin on localisation of laminin receptor, integrin α6 (Sonawane et al., 2005). Further clonal depletion of E-cadherin in periderm by injecting cdh1 MO shows an appreciable increase in membrane localisation of Lgl2 protein in juxtaposed cells in the basal epidermis and vice versa (Arora et al., 2020). I sought to understand if loss of Laminin α5, in addition to its effect on cell-cell junction protein, E-cadherin, also affects membrane Lgl2 levels in the same layer. I found that lama5 mutants show lower levels of membrane Lgl2 (Fig 3b, n=3, ****p<0.0001, Mann Whitney test) in basal epidermis in comparison to wild type siblings. Membrane Lgl2 also mimics reduced cell height measured by junctional E-cadherin seen in lama5 mutants, in comparison to wild type siblings (Fig 3c, n=3, ****p<0.0001, Mann Whitney test). Thus, loss of Laminin α5 affects both cell-cell junctional protein E-cadherin, and basolateral cell polarity protein Lgl2 and is required to maintain normal membrane levels of both in basal epidermis.

**Fig. 3.**
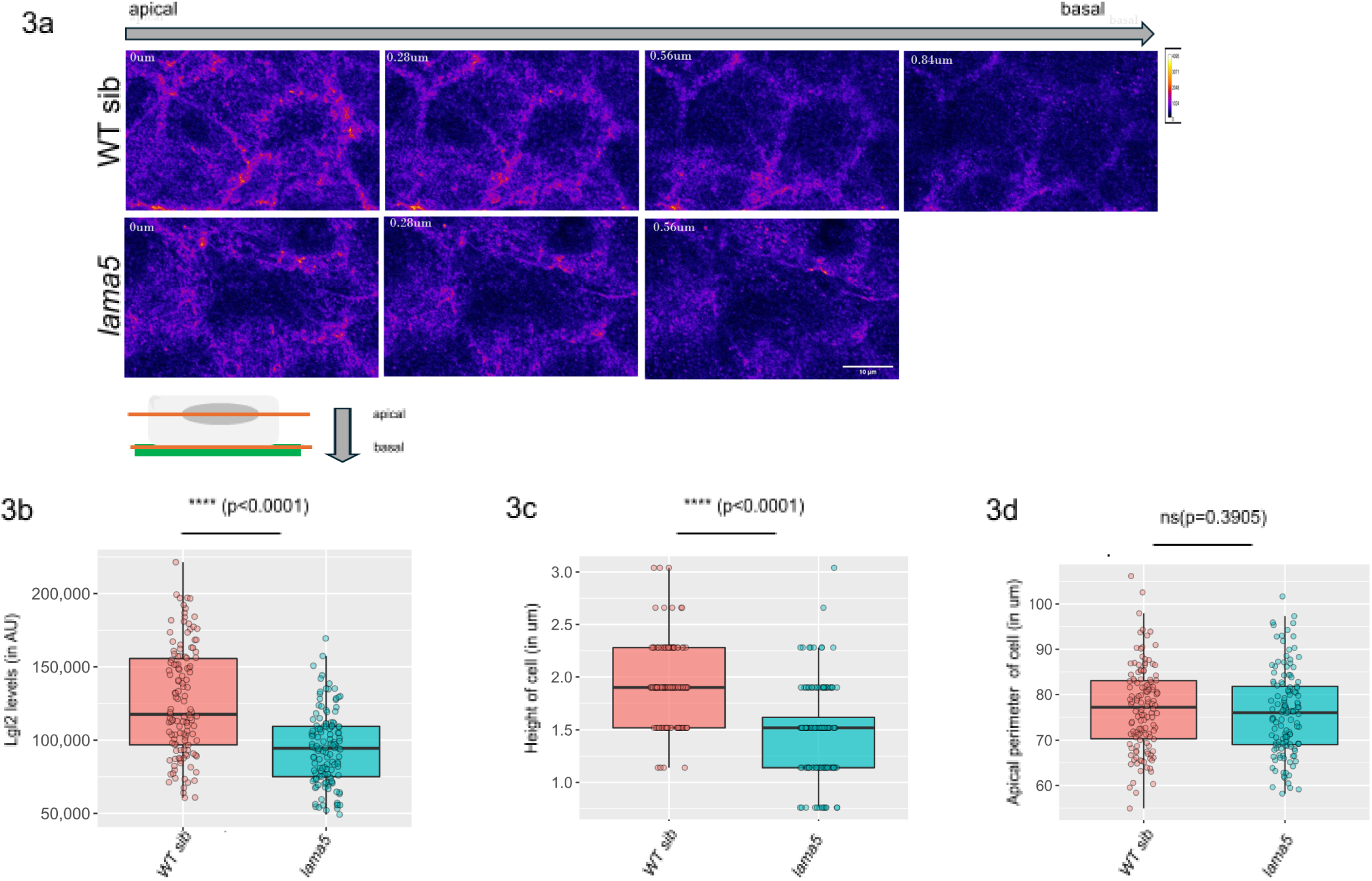
Basolateral membrane polarity protein Lgl2 level is affected in basal epidermis in *lama5* mutants and phenocopies cell morphology effects shown by junctional E-cadherin. Total membrane Lgl2 levels goes down in basal epidermis upon loss of laminin alpha5 (Fig3a, 3b) in basal epidermis. Cell height decreases in basal epidermis in *lama5* mutants (Fig3c). Apical perimeter does not change significantly in *lama5* mutants (Fig3d). Fig 3b, 3c ****p<0.0001 (n=3, Mann Whitney test); Fig 3d ‘ns’p=0.3905 means (non significant difference, n=3, unpaired t test). The schematic diagram depicts one basal epidermal cell resting on a basement membrane towards its basal side. ‘apical’ for basal epidermis lies above close to periderm. ‘basal’ for basal epidermis lies below towards the basement membrane region. (See supplementary S3, and S14 for source file and statistical comparisons). Scale bar depicts 10um length. Images are shown as heat map with calibration bar ranging from 0 to 4095 (in Fiji).

### 4. Basement membrane protein Laminin α5 is required to maintain cell-cell contact in basal epidermis via junctional E-cadherin

Loss of E-cadherin levels in cell-cell junction has been shown to be the hallmarks of cells undergoing epithelial-to-mesenchyme (EMT) transitions where they detach from each other (Hu et al., 2019; Paul et al., 2023; Ribatti et al., 2020; Sánchez-Tilló et al., 2010; Vesuna et al., 2008; Wong et al., 2014). I wanted to see if loss of junctional E-cadherin seen in lama5 mutants in basal epidermis show a similar effect of detachment seen during EMT. Live imaging CAAX-GFP injected embryos show that basal epidermal cells are detached from each other in lama5 mutants (Movie S2). Further these detached basal epidermal cells throw protrusions at cell boundaries. These protrusions were found to be highly dynamic over time showing retractions and extensions variably (Fig 4, green arrowhead in WT sib indicate intact cell-cell adhesion/boundary in comparison to detached cells and dynamic cell boundary protrusions seen in lama5 mutants at variable time points). The loss in cell shape (Fig 4, Movie S2), cell-cell detachment, and dynamicity seen in basal epidermis cells upon loss of Laminin α5 suggest that Laminin α5 plays role in maintaining epithelial characteristics of basal epidermal cells. In its absence, these cells tend to lose their epithelial nature, and show mesenchyme cell traits.

**Fig. 4.**
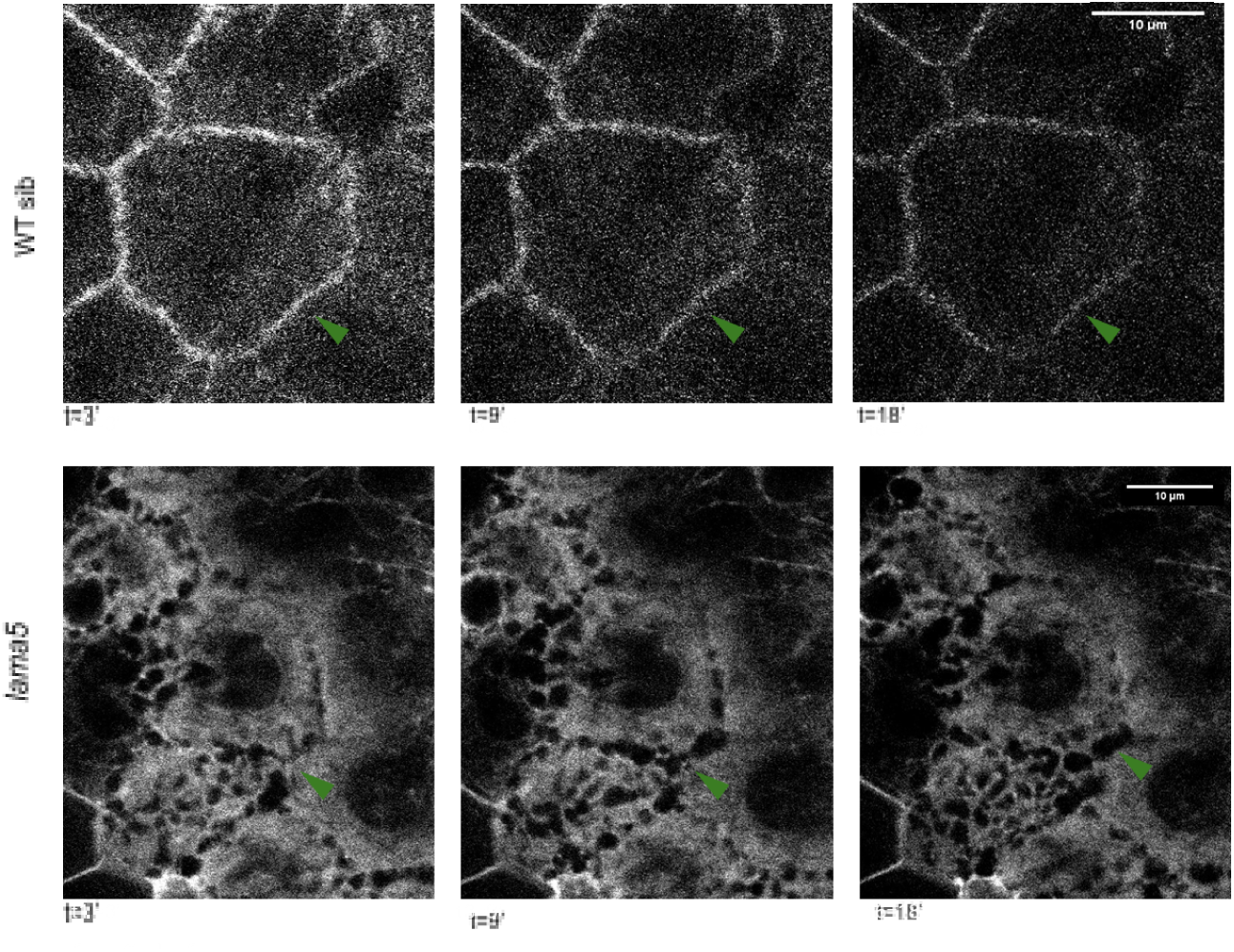
Loss of junctional E-cadherin in basal epidermis causes cell-cell detachment with cells throwing dynamic protrusions at cell boundaries. Live movie of CAAX-gfp injected embryos at 1-4 cell stage show intact cell shape and boundaries in *lama5* wt (T=13.44’) while in *lama5* mutants (T=21.12’) CAAX-gfp is diffuse filling up the cells that show cell-cell detachment and throw dynamic protrusions at cell periphery (Movie S4). Snapshots from live movies show the dynamic nature of cells in basal epidermis (Fig4). Green arrowheads indicate cell boundaries. Duration of live movie for *lama5* wt is 13.44’ (maximum intensity projection of few z-stacks z29-z36, no. of time frames=7, each time frame (T)=3.32’); duration of live movie for *lama5* mut is =21.12’ (single z-stack z44, no. of time frames=9, each time frame (T)=3.32’). Scale bar depicts 10um length. Images are shown in grays LUT (in Fiji).

### 5. Laminin α5 in the basement membrane is required to maintain epithelial proliferation of basal epidermal cells

Basal epidermal cells were found to lose their epithelial characteristics such as cell-cell adhesion, distinct cell boundaries, and epithelial cell shape showing more mesenchyme cell traits like loss of cell-cell adhesion, dynamic protrusions at cell boundaries, and variability in cell shape over time. This prompted us to look for other features characteristic of epithelial to mesenchyme transition (EMT) such as higher proliferation (Chen et al., 2020; Lee and Vasioukhin, 2008; Reischauer et al., 2009). I found that basal epidermis show an increase in proliferation marked by higher percentage of basal epidermal cells that are also positive for BrdU (Fig 5b, n=3, *p=0.0483, unpaired t test). Thus, in lama5 mutants, cells of basal epidermis show characteristic features of cell undergoing EMT which includes loss of junctional E-cadherin, detachment from neighbor cells, throwing dynamic protrusions at cell periphery, and increased proliferation in comparison to their wild type siblings in which basal epidermal cells maintain normal levels of junctional E-cadherin with intact cell-cell adhesion, cell shape, and baseline proliferation.

**Fig. 5.**
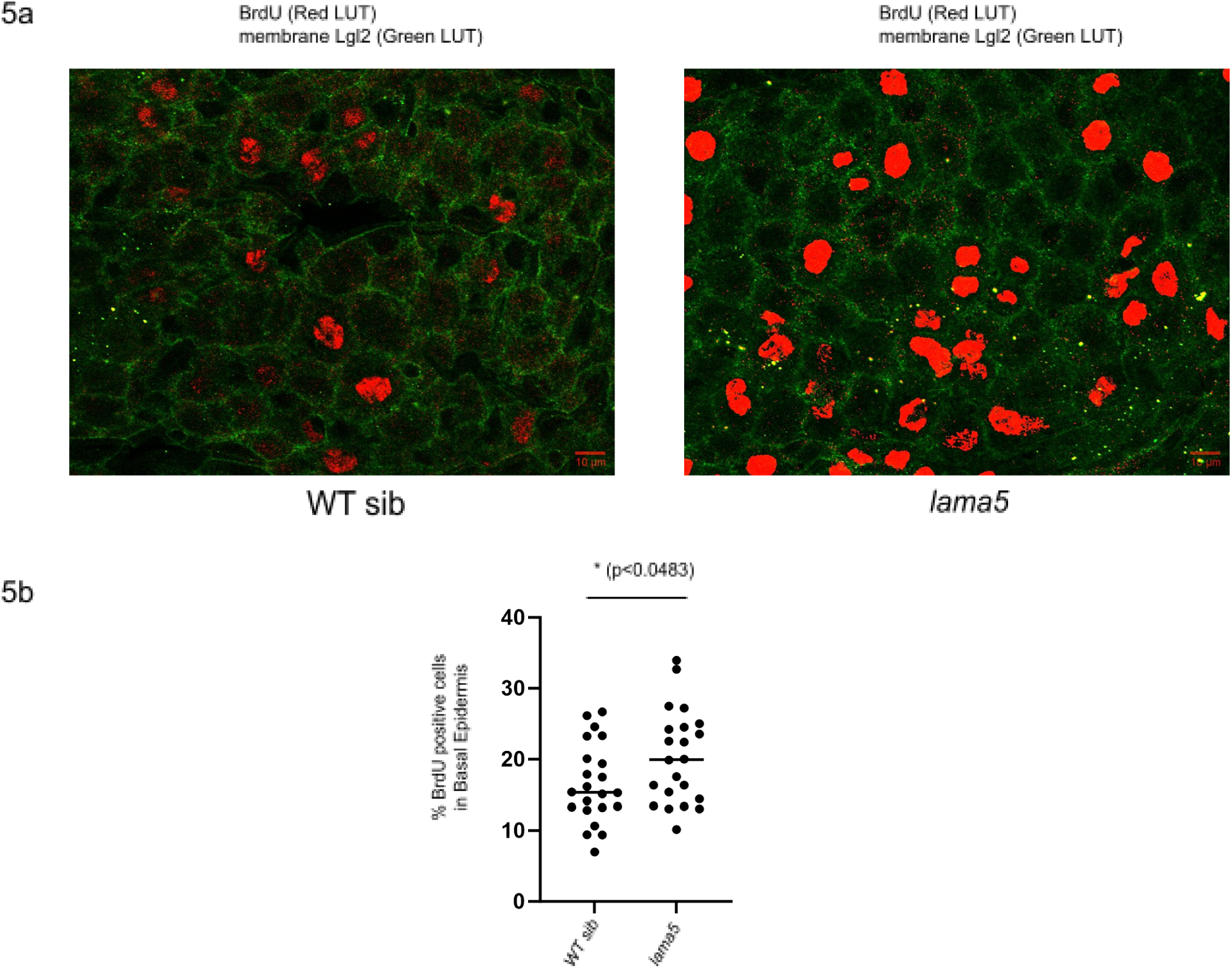
Basal epidermis shows higher proliferation marked by BrdU positive cells in *lama5* mutants. There is a significant increase in BrdU positive cells in basal epidermis of *lama5* mutants (Fig5a, 5b). (See supplementary S14 for source file and statistical comparisons) Scale bar depicts 10um length. In images, BrdU is shown in red LUT, and membrane Lgl2 in green LUT (in Fiji).

### 6. There is non-autonomous relay of cell-cell junctional cues via E-cadherin between periderm and basal epidermis

Zebrafish epidermis is two-layered during development (Arora et al., 2020; Chang and Hwang, 2011, Sonawane et al., 2009). I was curious to see if perturbation of cell polarity and cell junction in lower basal epidermis impacts the cell polarity and cell junction of the periderm layer above it. I found a reduction in E-cadherin localisation in periderm in lama5 mutants in comparison to wild type siblings (Fig 6b, n=3, ****p<0.0001, unpaired t test). This suggests a non-autonomous effect of loss of junctional E-cadherin in one layer (basal epidermis) on the other layer in contact (periderm). This is in concert with a previous study where clonal depletion of E-cadherin in basal epidermis was found to decrease junctional E-cadherin levels of juxtaposed cells in periderm and vice versa (Arora et al., 2020). In addition to lower E-cadherin levels at junctions, I found that periderm cell morphology was also affected in lama5 mutants with a reduction in cell height (Fig 6c, n=3, ****p<0.0001, Mann Whitney test) and an appreciable increase in apical perimeter (Fig 6d, n3, *p=0.0338, Mann Whitney test) in comparison to their wild type siblings. Thus, loss of junctional E-cadherin in basal epidermis seems to affect junctional E-cadherin levels of the periderm, and cell morphology layer non autonomously.

**Fig. 6.**
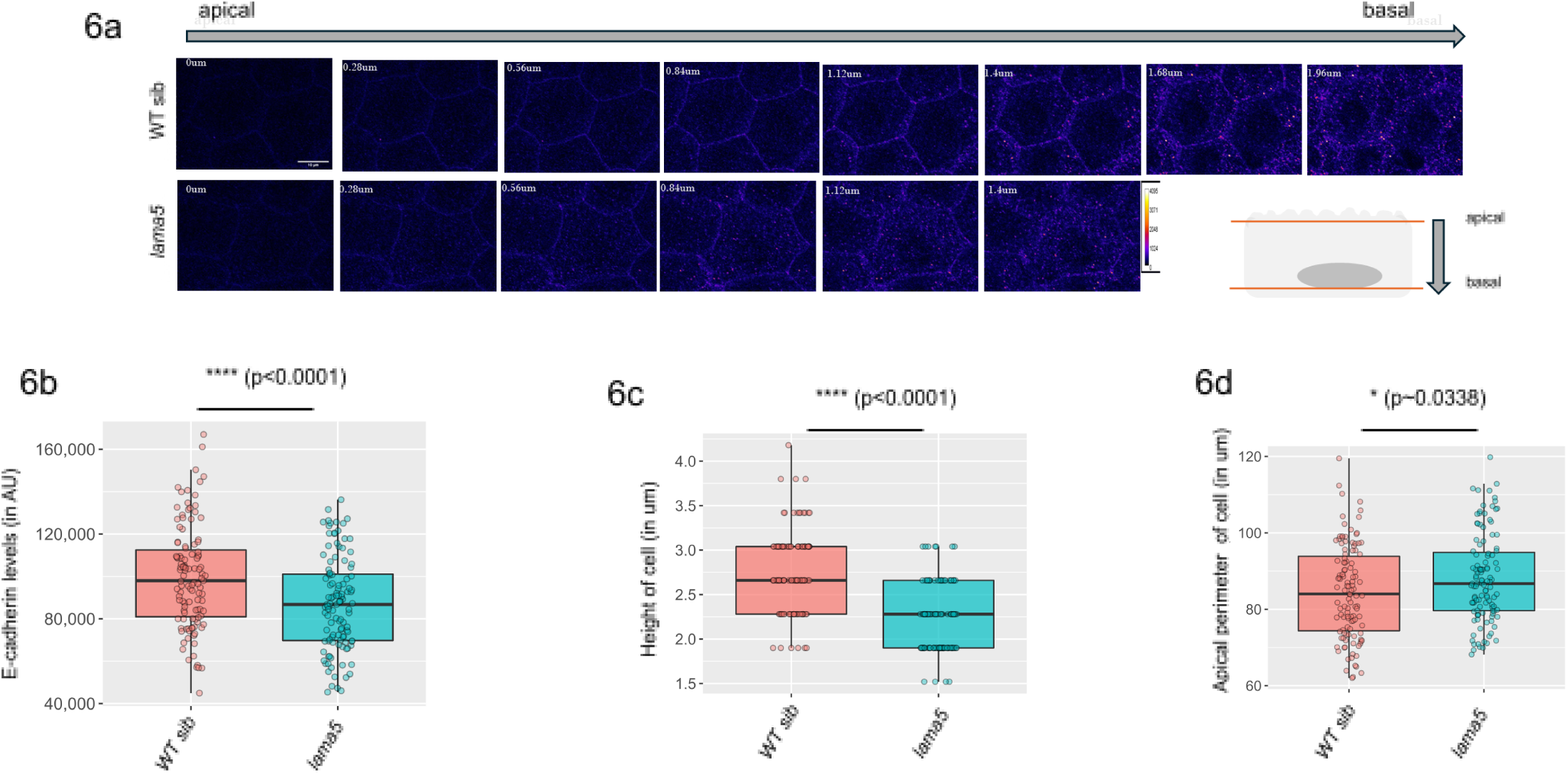
Junctional E-cadherin levels and cell morphology is affected in periderm in *lama5* mutants. Total junctional E-cadherin levels goes down in periderm upon loss of laminin alpha5 (Fig6a, 6b). Cell height decreases in basal epidermis in *lama5* mutants (Fig2c). Apical perimeter increases significantly in *lama5* mutants (Fig6d). Fig 6b, ****p<0.0001 (n=3, unpaired t test); Fig 6c ****p<0.0001 (n=3, Mann Whitney test); Fig 6d *p∼0.0338 (n=3, Mann Whitney test). The schematic diagram depicts one basal epidermal cell resting on a basement membrane towards its basal side. ‘apical’ for basal epidermis lies above close to periderm. ‘basal’ for basal epidermis lies below towards the basement membrane region. (See supplementary S6, and S14 for source file and statistical comparisons). Scale bar depicts 10um length. Images are shown as heat map with calibration bar ranging from 0 to 4095 (in Fiji).

### 7. Transduction of cell polarity and junction cues occurs between periderm and basal epidermis

Periderm cells have a distinct apicobasal cell polarity during development and later stages, where apical domain is marked by membrane aPKC, and F-actin based apical microridges while, Lgl2 is localised in its basolateral membranes (Arora et al., 2020; Sonawane et al. 2005, Sonawane et al., 2009, Pinto et al., 2019, Raman et al., 2016). In addition to lower junctional E-cadherin levels, I found that loss of Laminin α5 results in increased localization of membrane aPKC in its apical domain (Fig 8b, n=3, ****p=0.0001, Mann Whitney test; Fig 8c, n=3, ****p<0.0001, unpaired t test) in comparison to wild type siblings. This suggests a reinforced apical polarity seen in lama5 mutants by increasing apical membrane localisation of aPKC in order to maintain distinct apical domain characteristic of epithelial cells. I also checked the levels of membrane Lgl2 in periderm which did not show significant difference between lama5 mutants and their wild type siblings (Fig 7b, n=3, ‘ns’ p=0.1556, Mann Whitney test) suggesting that membrane basolateral domain identity is not perturbed upon loss of Laminin α5. Thus, despite the loss of cell polarity and junctions seen in basal epidermis in lama5 mutants, the periderm in lama5 mutants is able to maintain its unique apicobasal identity characteristic of epithelial cells by reinforcing membrane aPKC localisation apically and maintaining normal Lgl2 levels basolaterally. Further, the membrane Lgl2 show a similar decrease in cell height (Fig 7c, n=3, ****p<0.0001, Mann Whitney test) as shown by junctional E-cadherin in lama5 mutants in comparison to wild type siblings. However, apical perimeter shows a slight non-significant increase in lama5 mutants in comparison to their wild type siblings (Fig 7d, n=3, ‘ns’ p=0.1369, Mann Whitney test). Thus membrane Lgl2 seems to mimic changes in cell morphology of periderm seen in lama5 mutants in comparison to their wild type siblings as shown by junctional E-cadherin.

**Fig. 7.**
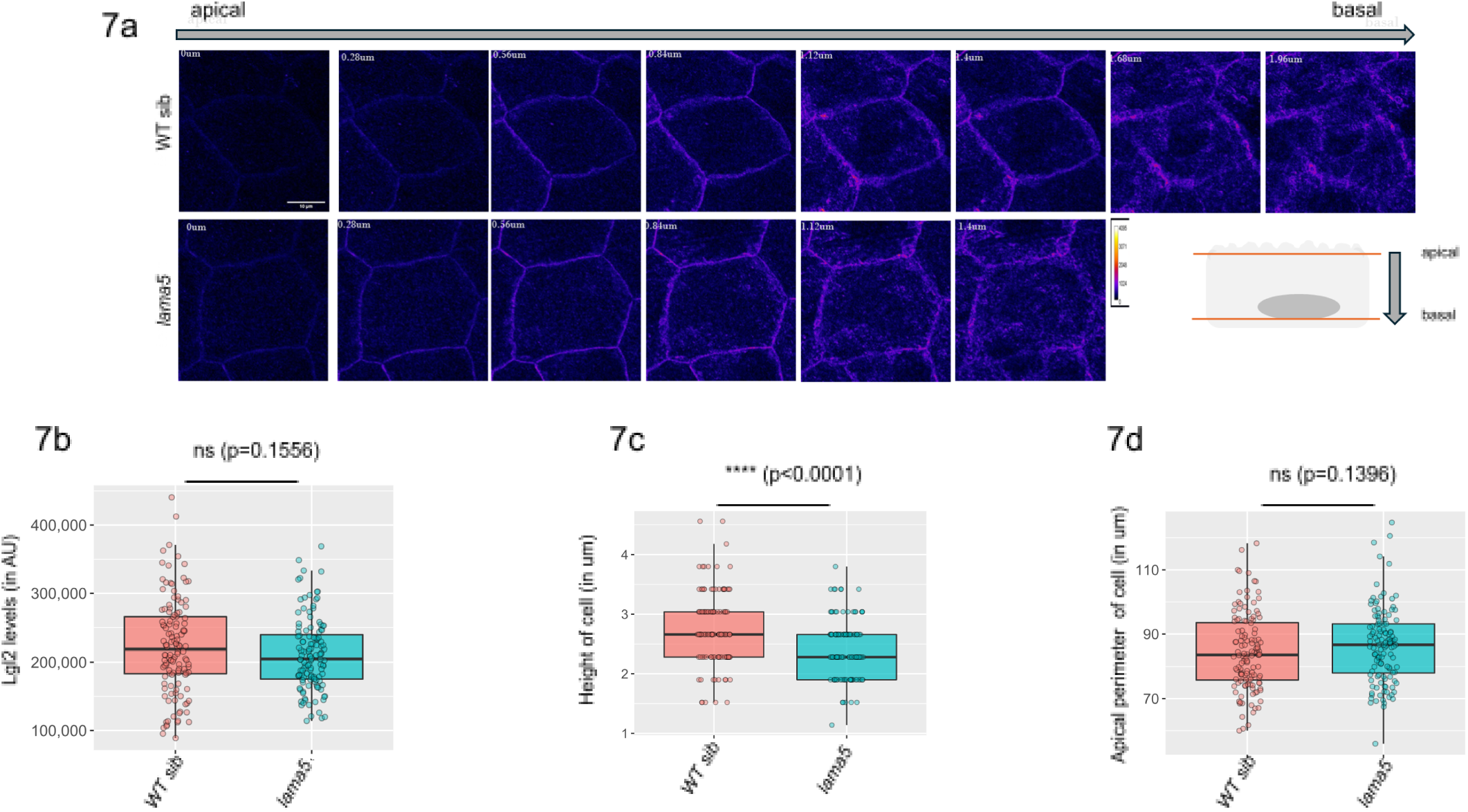
Basolateral membrane polarity protein Lgl2 level is not significantly affected in periderm in *lama5* mutants and phenocopies cell morphology effects shown by junctional E-cadherin. Total membrane Lgl2 level is unaffected in periderm upon loss of laminin alpha5 (Fig7a, 7b). Cell height decreases in periderm in *lama5* mutants (Fig7c). Apical perimeter does not change significantly in *lama5* mutants (Fig7d). Fig 7b ‘ns’ p=0.1556 means (non significant difference, n=3, Mann Whitney test); Fig 7c ****p<0.0001 (n=3, Mann Whitney test); Fig 7d ‘ns’ p=0.1396 means (non significant difference, n=3, Mann Whitney test). The schematic diagram depicts one basal epidermal cell resting on a basement membrane towards its basal side. ‘apical’ for basal epidermis lies above close to periderm. ‘basal’ for basal epidermis lies below towards the basement membrane region. (See supplementary S6, and S14 for source file and statistical comparisons). Scale bar depicts 10um length. Images are shown as heat map with calibration bar ranging from 0 to 4095 (in Fiji).

**Fig. 8.**
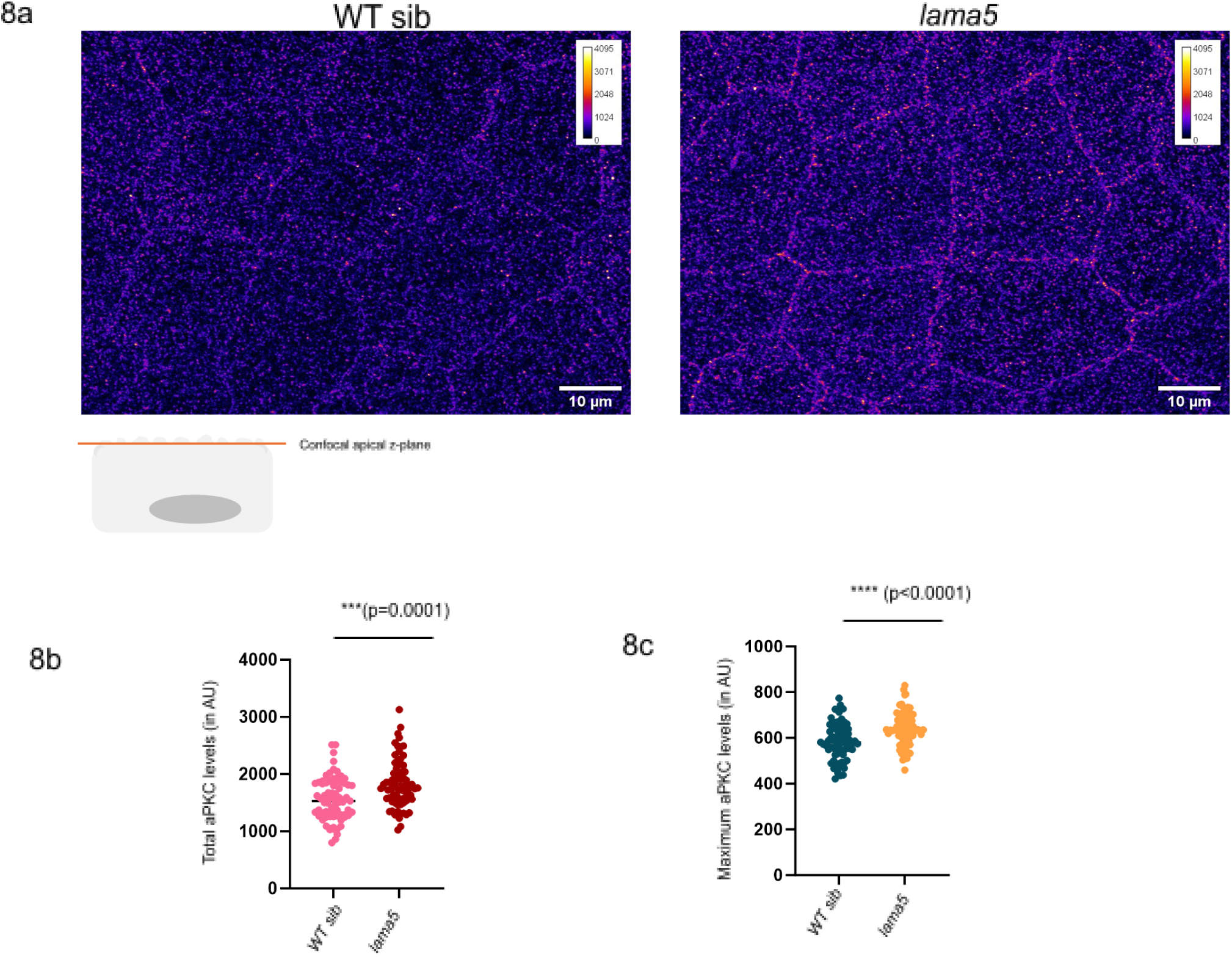
Apical membrane polarity protein aPKC level increases in periderm in *lama5* mutants. Total membrane aPKC (Fig 8b) level and maximum membrane aPKC (Fig8a, 8c) level significantly increased in periderm upon loss of laminin alpha5. Fig 8b ****p=0.0001 (n=3, Mann Whitney test); Fig 8c ****p<0.0001 (n=3, unpaired t test).The schematic diagram depicts one periderm cell. Confocal apical z-plane for periderm lies uppermost close to the microridge region. (See supplementary S14 for source file and statistical comparisons). Scale bar depicts 10um length. Images are shown as heat map with calibration bar ranging from 0 to 4095 (in Fiji).

### 8. aPKC levels are required to maintain microridge length and patterns

Previous studies (Raman et al, 2016) have shown that aPKC loss seen in has/aPKC mutants result in longer microridge length. Conversely, overexpression of aPKC in clonal fashion shows shorter microridges(Sen et al., 2024). I sought to understand the effect of increase in aPKC levels endogenously upon microridges which has not been addressed previously. I found that higher levels of endogenous aPKC found in lama5 mutants result in shorter microridge length in apical domains (Fig 9b, n=3, **p=0.0010, Mann Whitney test) in comparison to the wild type siblings. These short microridges found in lama5 mutants were more homogenous in length showing lower standard deviation in microridge length in comparison to their wild type siblings (Fig 9c, n=3, **p=0.0063, Mann Whitney test). Microridges form complex F-actin based patterns with intersections or branch points seen in wild type cells. I found a reduction in number of branch points or intersections in lama5 mutants in comparison to their wild type siblings (Fig 9e, n=3, **p=0.0036, Mann Whitney test).Thus lama5 mutants have shorter microridge length and form less complex F-actin based patterns. Further, the area fraction, i.e., percentage area of cell covered by microridges was significantly reduced in lama5 mutants in comparison to their wild type siblings (Fig 9d, n=3, **p=0.0012, unpaired t test). The total number of microridges did not show significant change between lama5 mutants and its wild type siblings (Fig 9f, n=3, ‘ns’ p=0.6731, Mann Whitney test). These results suggest that lama5 mutants have shorter microridge length, form less complex patterns, and cover lesser cell area in comparison to their wild type counterparts which might be attributed to higher levels of membrane aPKC found apically in lama5 mutants in comparison to wild type siblings.

**Fig. 9.**
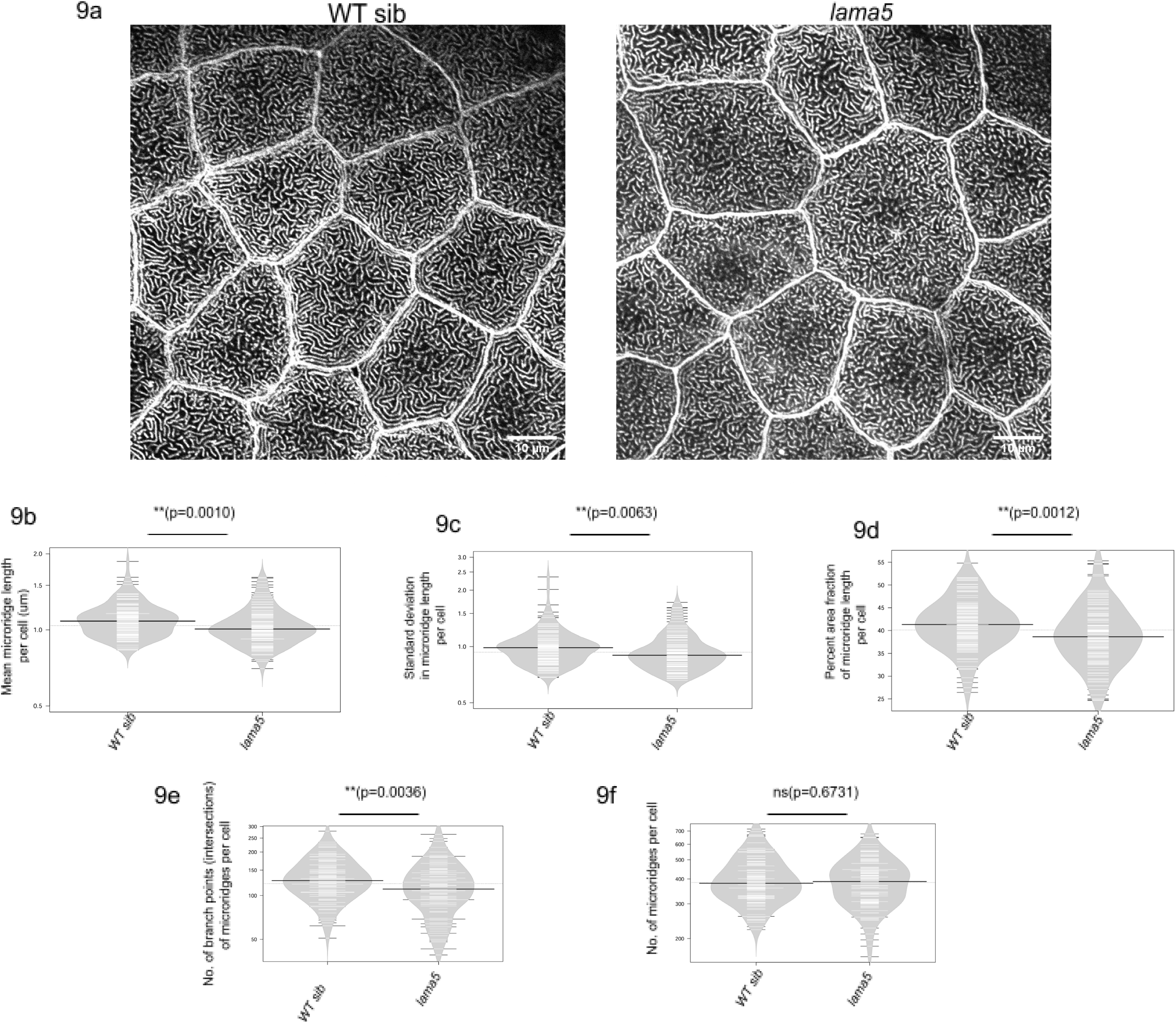
F-actin based apical polarity structures microridges in periderm form smaller and less complex patterns in *lama5* mutants. Mean microridge length (Fig 9a, 9b), standard deviation in mean microridge length (Fig 9c), percent area fraction covered by microridges per cell (Fig 9d), no. of branch points or intersections formed by microridges per cell (Fig 9e) decreases in *lama5* mutants. Total no. of microridges per cell does not show significant difference in *lama5* mutants (Fig 9f). Fig 9b **p=0.010, Fig9c **p=0.0063, Fig9d **p=0.0012, **p=0.0036 (n=3, Mann Whitney test); Fig 9f ‘ns’ p=0.6731 means (non significant difference, n=3, Mann Whitney test). (See supplementary S9, and S14 for source file and statistical comparisons). Scale bar depicts 10um length. Images are shown in grays LUT (in Fiji).

### 9. Periderm is able to maintain its cell-cell adhesion despite decrease in junctional E-cadherin levels independently

The reduction in junctional E-cadherin levels prompted us to see if periderm is showing any traits of EMT as evident in basal epidermis. I found that CAAX-GFP localised to the cell membranes of periderm cells in both lama5 mutants and wild type siblings. Periderm cells in lama5 mutants were able to maintain cell-cell adhesion and intact cell shape like periderm cells in wild type siblings over time (Movie S3, Fig 10, green arrowheads in WT sib and lama5 mutants indicate intact cell-cell adhesion/boundary at variable time points). Thus, in a two-layered epidermis found in zebrafish during development, it seems that despite one layer (basal epidermis) undergoing EMT by losing its epithelial integrity and gaining more mesenchymal traits, the other layer (periderm) is able to maintain its cell-cell adhesion and cell shape characteristic of epithelial tissue.

**Fig. 10.**
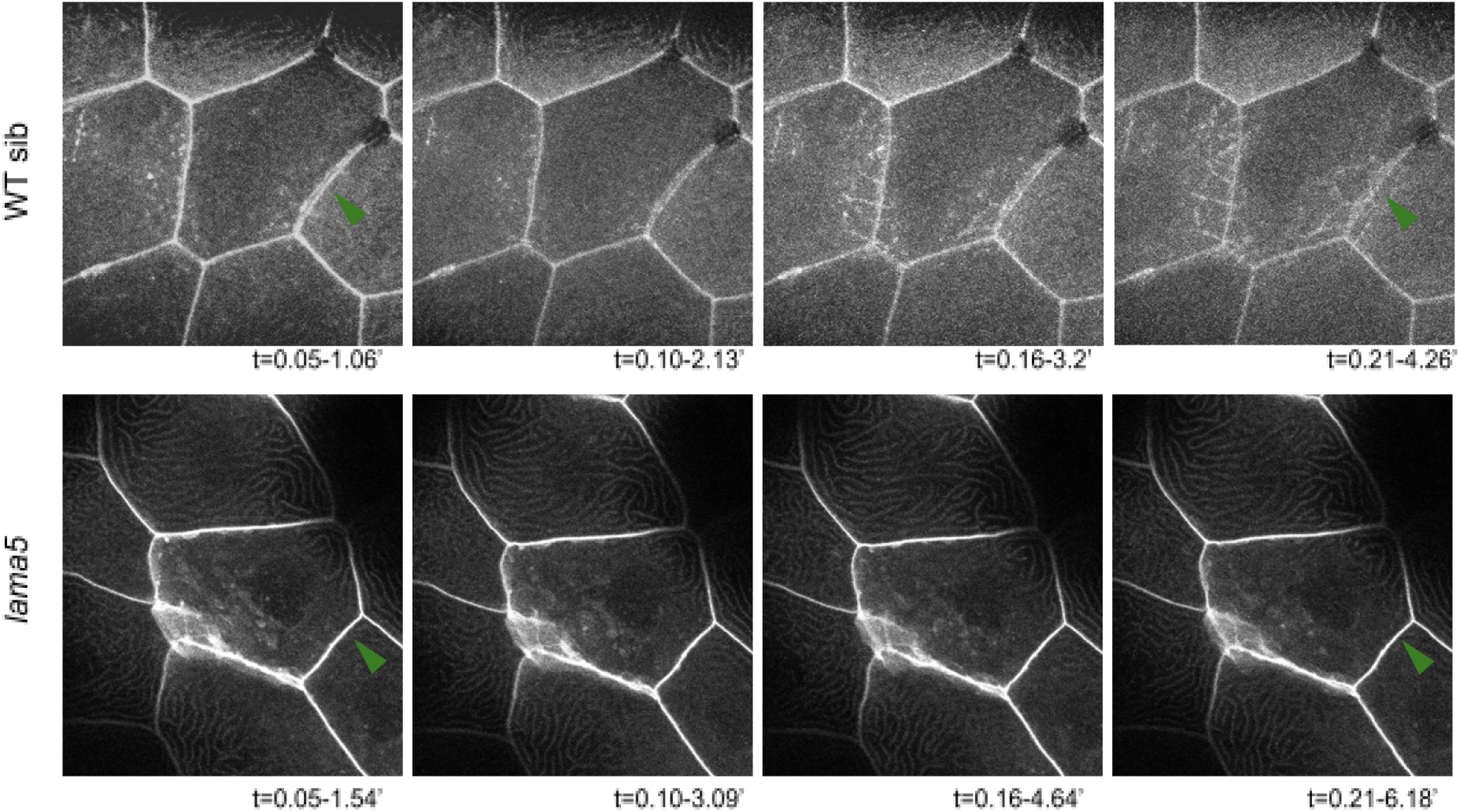
Periderm shows intact cell-cell adhesion and cell shape in *lama5* mutants. Live movie of CAAX-gfp injected embryos at 1-4 cell stage show intact cell shape and boundaries in *lama5* wt and *lama5* mutants (Movie 10). Snapshots from live movies show intact cell boundaries and cell shape in the periderm of *lama5* wt and *lama5* mutants (Fig10). Green arrowheads indicate cell boundaries. Duration of live movie for *lama5* wt is 8.53’ (maximum intensity projection of few z-stacks z1-z19, no. of time frames=8, each time frame (T)=3.32’); duration of live movie for *lama5* mut is 12.37’ (maximum intensity projection of few z-stacks z1-z29, no. of time frames=8, each time frame (T)=3.32’). Scale bar depicts 10um length. Images are shown in grays LUT (in Fiji).

### 10. Laminin α5 acts via Integrin α6b receptor to maintain epithelial characters of basal epidermis

Since laminin is found towards the basal side of basal epidermis in the basement membrane region, I wanted to ask how it affects epithelial junction and polarity molecules of basal epidermis. Integrins are heterodimeric cell surface receptors formed of α and β subunits, which localize to the basal domain and mediate cell-matrix adhesion by interacting with laminins (Arimori et al., 2021; Khademi et al., 2023; Liu et. al., 2023; Nishiuchi et al., 2006; Zhang et al., 2019). I found that itga6b was highly enriched in zebrafish epidermis during development (Danio cell dataset; Farrell et al., 2018; Sur et al., 2023). Further, I found that itga6b mutants like lama5 mutants also show reduction in junctional E-cadherin levels in basal epidermis (Fig 11b, n=3, *p=0.0097, Kruskal Wallis test, Dunn’s post hoc test) in comparison to wild type siblings. Further, itga6b mutants show reduction in cell height in basal epidermis (Fig 11c, n=3, *p=0.0196, Kruskal Wallis test, Dunn’s post hoc test) in comparison to wild type siblings. The apical perimeter was however found to increase in itga6b mutants in comparison to lama5 mutants (Fig 11d **p=0.0033, n=3, Kruskal Wallis test, Dunn’s post hoc test). Thus, it appears that Laminin α5 in the basement membrane region binds to Integrin α6b receptors to maintain its epithelial cell-cell adhesion and morphology.

**Fig. 11.**
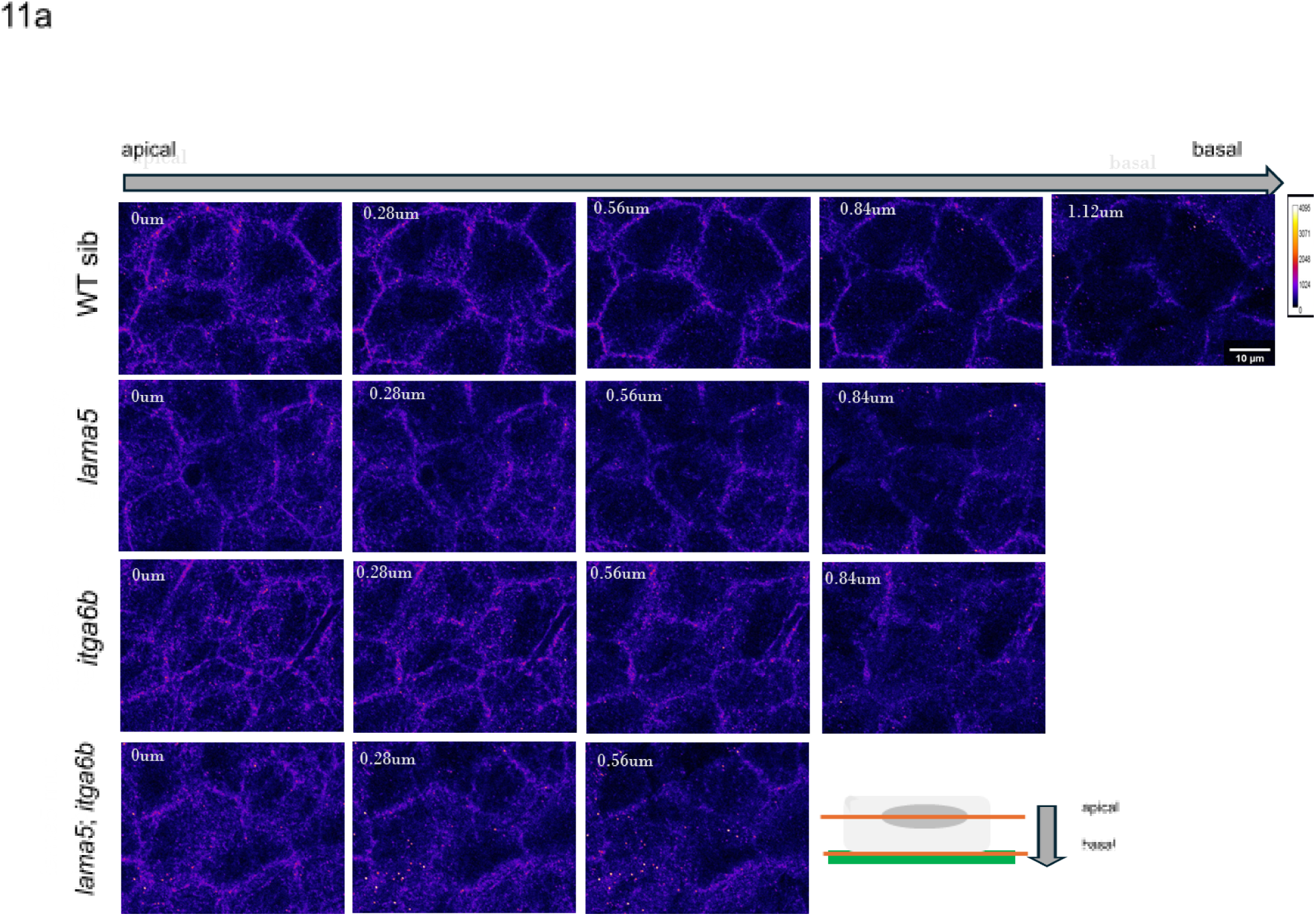
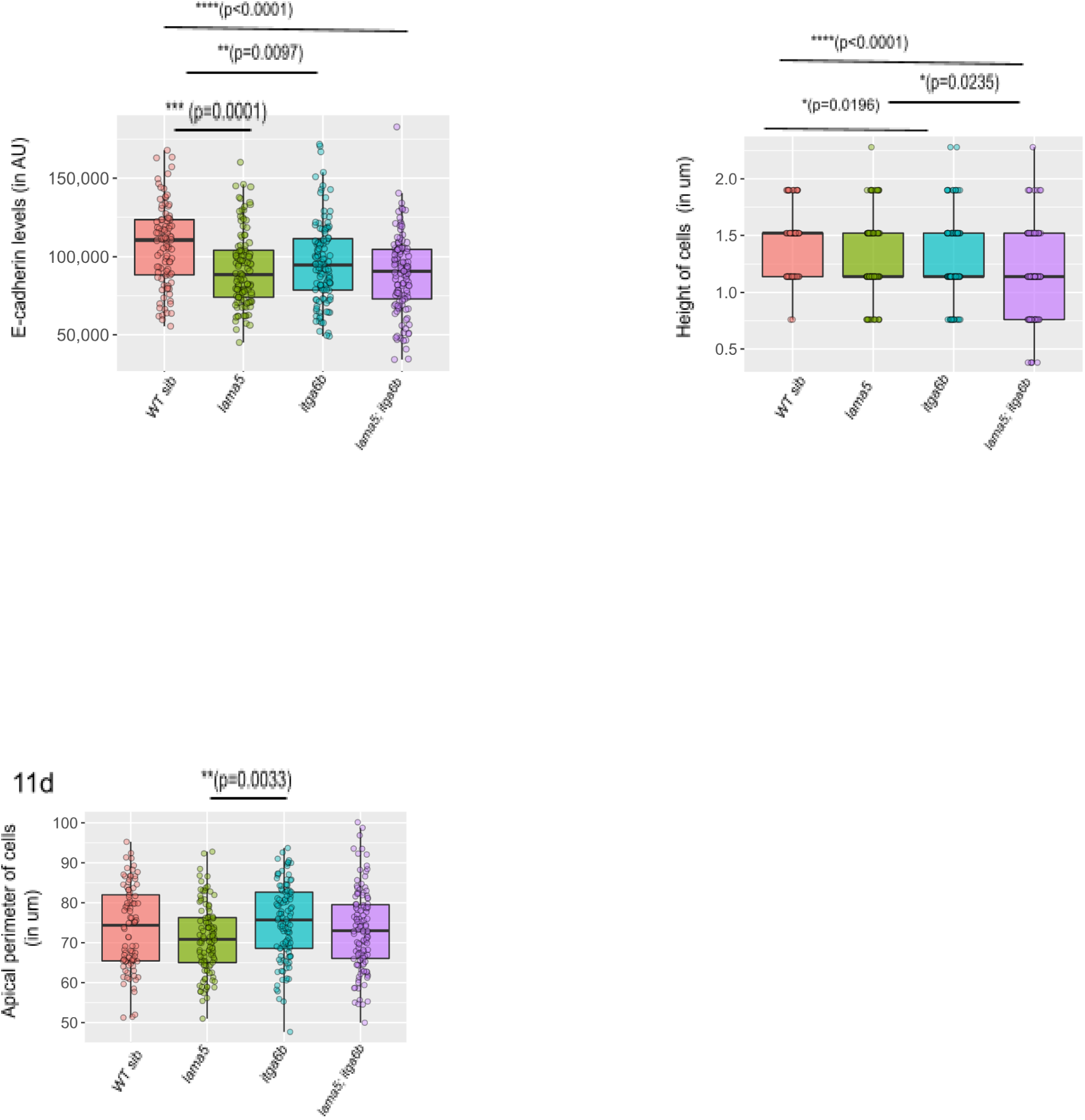
Junctional E-cadherin levels and cell morphology is affected in basal epidermis in *itga6b* and *lama5;itga6b* like *lama5*. Total junctional E-cadherin levels goes down in basal epidermis upon loss of *itga6b* and *lama5;itga6b* as seen in *lama5* (Fig11a, 11b). Cell height decreases in basal epidermis in *lama5* and *itga6b* which is further exacerbated in *lama5;itga6b* (Fig11c). Apical perimeter does not change significantly except between *lama5* and *itga6b* (Fig11d). Fig 11b, ***p=0.0001 (wt vs *lama5*, n=3, Kruskal Wallis test, Dunn’s post hoc test), **p=0.0097 (wt vs *itga6b*, n=3, Kruskal Wallis test, Dunn’s post hoc test), ****p<0.0001 (wt vs *lama5;itga6b*, n=3, Kruskal Wallis test, Dunn’s post hoc test); Fig 11c *p=0.0196 (wt vs *itga6b*, n=3, Kruskal Wallis test, Dunn’s post hoc test), ****p<0.0001 (wt vs *lama5;itga6b*, n=3, Kruskal Wallis test, Dunn’s post hoc test), **p=0.0235 (*lama5* vs *lama5;itga6b*, n=3, Kruskal Wallis test, Dunn’s post hoc test); Fig 11d **p=0.0033 (*lama5* vs *itga6b*, n=3, Kruskal Wallis test, Dunn’s post hoc test). The schematic diagram depicts one basal epidermal cell resting on a basement membrane towards its basal side. ‘apical’ for basal epidermis lies above close to periderm. ‘basal’ for basal epidermis lies below towards the basement membrane region. (See supplementary S11, and S14 for source file and statistical comparisons). Scale bar depicts 10um length. Images are shown as heat map with calibration bar ranging from 0 to 4095 (in Fiji).

**Fig. 12.**
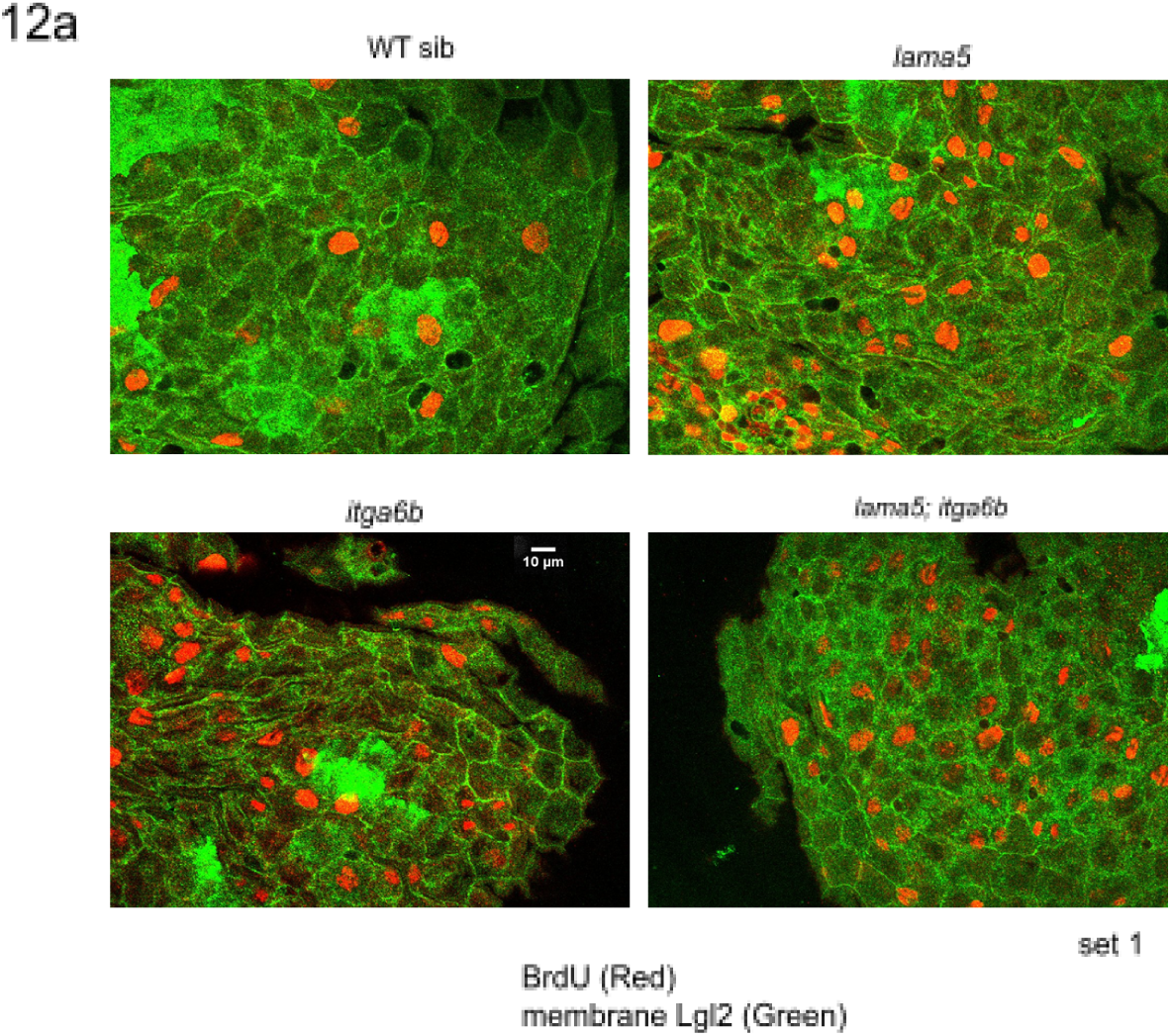
Basal epidermis shows higher proliferation marked by BrdU positive cells in *lama5* and/or *itga6b* mutants. There is an increase in BrdU positive cells in basal epidermis of *lama5* and/or *itga6b* mutants seen across two sets. (Fig 12, Fig S12). Scale bar depicts 10um length. In images, BrdU is shown in red LUT, and membrane Lgl2 in green LUT (in Fiji)..

### 11. Laminin α5 and Integrin α6b act in a single pathway to maintain epithelial cell-cell adhesion

I wanted to understand if Laminin α5 and Integrin α6b bind to other receptors and ligands respectively in order to maintain epithelial cell-cell adhesion in basal epidermis. However, loss of both Laminin α5 and Integrin α6b seen in lama5; itga6b double mutants did not worsen the cell-cell adhesion phenotype and show a similar reduction in junctional E-cadherin levels in comparison to wild type siblings (Fig 11b, n=3, ***p<0.0001, Kruskal Wallis test, Dunn’s post hoc test) like that seen in lama5, and itga6b single mutants. Thus, it appears that Laminin α5 via Integrin α6b solely is required to maintain epithelial cell-cell adhesion in basal epidermis. However, there is an exacerbation in cell height phenotype where loss of both Laminin α5 and Integrin α6b in lama5; itga6b double mutants do not show a further significant decrease in cell height in comparison to lama5 and itga6b single mutants (Fig 11c****p<0.0001 (wt vs lama5;itga6b), n=3, Kruskal Wallis test, Dunn’s post hoc test), **p=0.0235 (lama5 vs lama5;itga6b), n=3, Kruskal Wallis test, Dunn’s post hoc test). This might be due to other pathways that impinge on the actin cytoskeleton via Laminin α5 and/or Integrin α6b independently, to maintain epithelial cell morphology in basal epidermis (Akhtar and Streuli, 2013; Cohen et al., 2011; Raman et al, 2018, Roignot et al., 2013).

### 12. Laminin α5 acts via Integrin α6b to maintain epithelial characteristics of basal epidermis

Since loss of Laminin α5 and/or Integrin α6b show similar reduction in junctional E-cadherin levels, I went on to see for other characteristics that define epithelial nature of basal epidermis in itga6b single mutants, and lama5;itga6b double mutants. I found that either loss of Integrin α6b alone, or both Laminin α5 and Integrin α6b, show similar phenotype like that seen upon loss of Laminin α5 alone, viz., basal epidermal cells detach from each other showing dynamic protrusions at cell periphery (Fig 13a, green arrowhead in WT sib indicate intact cell-cell adhesion/boundary in comparison to detached cells and dynamic cell boundary protrusions seen upon loss of lama5 and/or itga6b alleles at variable time points). Interestingly, this phenotype was also seen upon loss of a single allele of lama5 and/or itga6b. Thus, Laminin α5 ligand binds to Integrin α6b receptor to maintain epithelial cell-cell adhesion and epithelial cell shape of basal epidermal cells during development. In the absence of such ligand-receptor binding, the basal epidermal cells tend to lose their epithelial nature and transition towards mesenchyme-like cell state, showing dynamic cell boundaries and loss of cell-cell adhesion, seen during EMT.

**Fig. 13.**
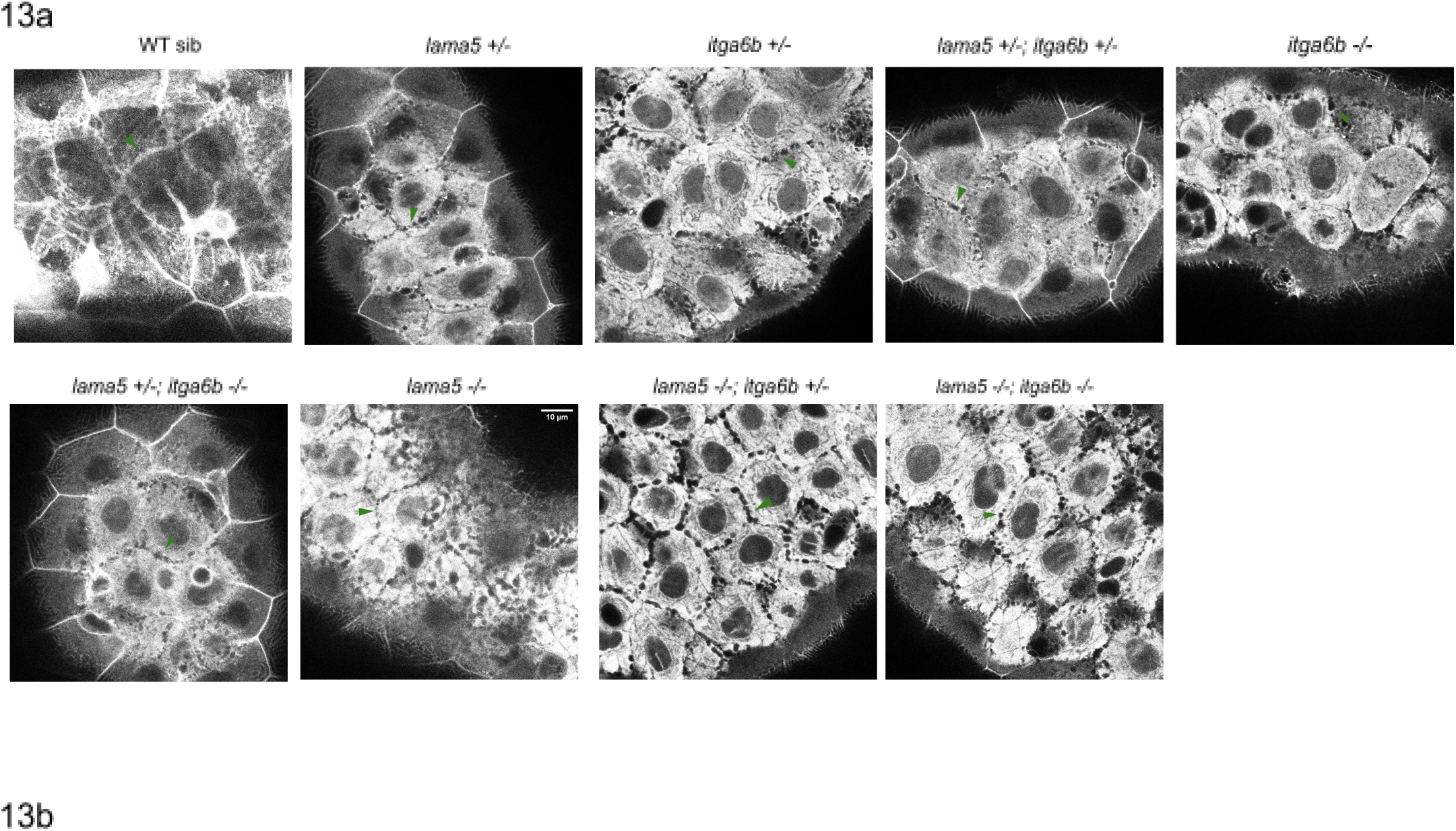
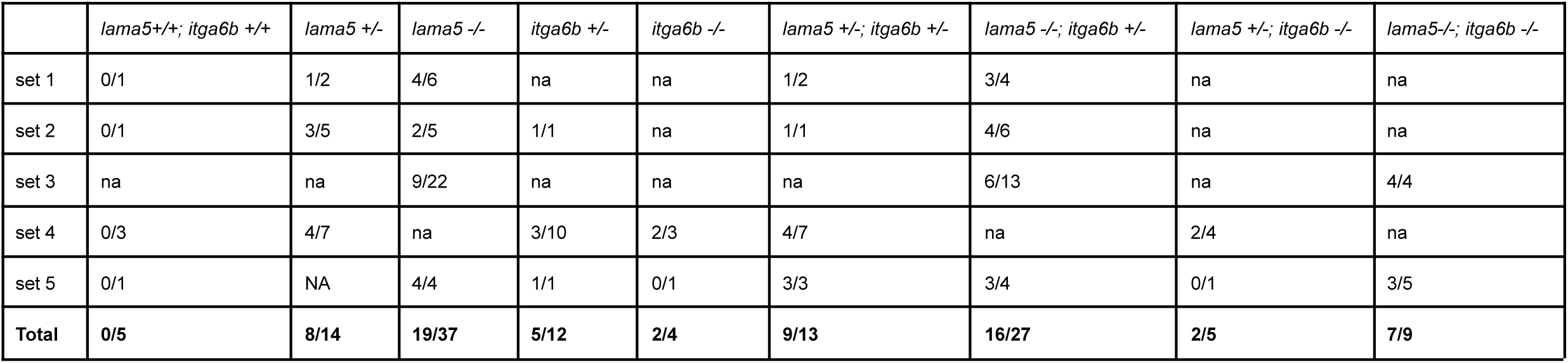
Loss of junctional E-cadherin in basal epidermis causes cell-cell detachment with cells throwing dynamic protrusions at cell boundaries upon single and/or double loss of *lama5* and/or *itga6b* allele. Live snapshots of CAAX-gfp injected embryos at 1-4 cell stage show intact cell shape and boundaries in wt while in *lama5* and/or *itga6b* single/double allele mutants show cell-cell detachment and dynamic protrusions at cell periphery (Fig13a). Table (Fig13b) shows no. of embryos that show this phenotype. Green arrowheads indicate cell boundaries. ‘na’ means ‘not available’. (See supplementary Fig. S13 for *lama5* and *itga6b* allele). Scale bar depicts 10um length. Images are shown in grays LUT (in Fiji).

## Discussion

Laminin proteins have long been recognised as basement membrane markers enriched towards the basal side of cells (Cohen et al., 2011; Matlin et al., 2017). They are known to sculpt tissue shape and structure during morphogenesis and development (Hamill et al., 2011, Miner et al., 1998; Webb et al., 2007; Yurchenco et al., 2004). Despite their enrichment only towards the basal side of the cell in close apposition to the membrane, it is a highly underappreciated polarity molecule in the field. In this study, I find that laminin molecules enriched only in the basement membrane region are required to maintain key epithelial characteristics of basal epidermal cells. Loss of laminin was found to compromise the normal localisation of E-cadherin at cell-cell junctions. Localisation of E-cadherin at cell-cell junctions is highly important for maintaining cell-cell adhesion which gives epidermis its unique epithelial identity showcased by intact cell junctions and cell shape. Upon loss of Laminin α5 many of these features such as intact cell-cell adhesion and cell shape were compromised. Live imaging of lama5 mutants show that in the absence of Laminin α5 in basement membrane region the basal epidermal cells tend to lose their epithelial features found in wild type siblings and acquire more mesenchymal traits such dynamic cell protrusions at cell boundaries, and detachment of cells from each other. This acquisition of dynamic cell feature along with loss of cell-cell adhesion is highly characteristic of cells undergoing epithelial to mesenchymal transition (EMT). Other features of cells undergoing EMT such as increased proliferation was also seen in lama5 mutants in comparison to their wildtype counterparts. Thus, it becomes evident that Laminin α5 enriched in the basement membrane region binds to receptors to maintain epithelial cell nature of basal epidermal cells in the absence of which cells lose their epithelial features and transition towards mesenchyme cell state.

Laminin proteins are present outside the cell in basement membrane region closely apposed to basal cell membrane.This basal cell membrane is highly enriched in receptors such as integrins, dystroglycans, and other receptors (Hamill, 2011, Nonnast et al., 2025; Yurchenco, 2011). I was interested to see which of the laminin receptors bind to it to mediate the epithelial cell state of basal epidermal cells. I found that loss Integrin α6b, a highly enriched Integrin isoform in epidermis, phenocopy loss of Laminin α5 showing reduction in junctional E-cadherin levels, and reduced cell height in basal epidermis. It also shows other phenotypes seen in lama5 mutants such as cell-cell detachment and increased cell dynamicity. Thus, it becomes clear that Laminin α5 binds to Integrin α6b receptor on the basal side of cells in basal epidermis in order to maintain its epithelial characteristics and prevent the cells from undergoing EMT. To my surprise, double loss of both Laminin α5 and Integrin α6b did not show exacerbation in these phenotypes except for reduction in cell height which is addressed later. Thus, it suggests that Laminin α5 acts solely via Integrin 6b pathway to maintain epithelial integrity of basal epidermal cells in the absence of which cells lose their epithelial structure and show features of EMT.

All studies so far on Laminins and cell polarity have been done in 2D & 3D-organoid cell culture systems, Drosophila follicle & gut epithelium, or mouse embryonic epidermis that consist of a single organised cell layer (Chen et al., 2018; Guo et al, 2024; Lerner et al., 2010; Miner et al,1998; Yu et al, 2005). Since zebrafish epidermis is organised in a bilayered fashion during development, I took advantage of it and went on to see the effect of loss of epithelial integrity in basal epidermis on the layer above it called periderm. Previous studies have hinted that there might be some non-autonomous crosstalks between periderm and basal epidermis (Arora et al., 2020). Here, I thoroughly characterised the effect of loss of cell-cell adhesion and cell polarity in one layer on cell polarity and junctions of other juxtaposed layer in lama5 mutants. I show that in lama5 mutants, periderm shows reduction in junctional E-cadherin levels due to layer non-autonomous effect of reduced junctional E-cadherin levels seen in basal epidermis. Despite this reduction, periderm is still able to maintain its cell-cell adhesion in lama5 mutants similar to that seen in wild type siblings. Further, periderm was successful in maintaining its unique apicobasal identity by increased apical localisation of membrane aPKC and maintaining similar wild type levels of membrane Lgl2 basolaterally. Thus, in a two-layer epidermal tissue I found that loss of cell-cell adhesion and polarity in one layer does not seem to compromise the cell-cell adhesion and apicobasal cell polarity of the other layer in contact with it. This result is highly interesting as it hints towards a survival benefit imposed upon a multilayered tissue in comparison to singly-layered epidermis. Here I show for the first time that despite loss of epithelial integrity in one layer, the other layer maintains its epithelial nature and shape to prevent the entire epidermal tissue from falling apart. Whether this surveillance mechanism imposed on bilayered tissue is unique to zebrafish or can be extended to other multilayered epidermal tissue needs to be examined.

F-actin is one of the earliest regulators of polarity where several studies have shown it to be upstream in establishment and maintenance of different forms of cell polarity such as apicobasal, anterior posterior etc. and cell junctions (Koch and Rose, 2023; Raman et al., 2018;, Roignot et al; 2013). I found that F-actin based microbridges were short and formed less complex patterns in periderm in lama5 mutants, suggesting an effect on F-actin based networks. This may be due to increase in apical membrane aPKC which is in concert with aPKC gain and loss of function studies that have shown its effect on microridge length (Raman et al., 2016, Sen et al., 2024). Further, I have also seen a reduction in cell height in both periderm and basal epidermis in lama5 mutants. I think that loss of epithelial cell-cell adhesion seen via detachment of cells from each other in basal epidermis in lama5 or itga6b mutants results in collapse in cell height which was found to exacerbate in lama5;itga6b double mutants. The exacerbated reduction in cell height seen in lama5;itga6b double mutants might be due to its more severe effect on actin cytoskeleton due to perturbation of additional pathways (Hamill et al., 2011; Raman et al., 2018; Roignot et al., 2013) that impinge on actin cytoskeleton via Rac1 upon double loss of both Laminin α5 and Integrin α6b. Further, we cannot rule out the bigger picture where collapse in cell height of basal epidermis may have a non autonomous effect on actin cytoskeleton of periderm resulting in reduced cell height and compensatory increase in apical perimeter of periderm cells in order to maintain epithelial homeostasis (Sonal et al., 2014; Lewis et al, 2020; Chouhan et al., 2024). Larger cell size can also be a contributing factor to shorter microridges due to less membrane available to form complex labyrinth like patterns (Lam et al., 2015; Pinto et al. 2019). However, how these factors impact microridge length and patterns remain to be elucidated.

Thus, my study for the first time show the importance of Laminin α5 in basement membrane region in maintaining epithelial identity of basal epidermal cells in the absence of which Laminin α5/Integrin α6b pathway is perturbed and cells lose their epithelial state and transition towards mesenchymal cell state showing features of epithelial to mesenchyme transition (EMT) in basal epidermis. I further elaborated how a two-layered tissue provides survival benefit over one layered epithelial tissue where despite the loss of epithelial integrity of one layer, the other layer is able to maintain its epithelial nature to prevent whole epidermal tissue from collapsing and promoting survival of the organism.

## Methodology and methods

### Fish strains

The fish lines used in this study were lama5tc17 (Carney et al., 2010), TgBAC(lamc1:lamc1-sfGFPp1) (Yamaguchi et al, 2022) and an unpublished Itga6b (fu36) allele obtained from Dr. Chaitanya Dingare and Prof. Virginie Lecaudey (Institute for Cell Biology and Neuroscience, Goethe University, Germany). The phenotype of lama5tc17 mutant at 42hpf was identified based on tail fin, pectoral fin, and yolk extension defects as shown in brightfield images below. Itga6bfu3 do not show any morphological defects during early development and were identified by genotyping. All experiments were done at 42hpf unless mentioned otherwise (Fig 1). BrdU pulse was given at 42hpf for 3hrs. Live imaging was performed from 1.8dpf-3dpf. For zebrafish maintenance and experimentation, the guidelines recommended by CPCSEA, Government of India, were followed.

### Genomic DNA extraction and PCR

Genomic DNA was extracted using hot-shot method. Whole embryo or its part was incubated in 30uL of 50mM NaOH for 15min at 95°C, vortexed, incubated on ice for 10min, 1/10 Tris-Cl(1M, pH8) was added, centrifuged at maximum speed for 10min and supernatant was used for PCR. Both laminin and integrin mutant alleles were identified by standard PCR. The reaction conditions were as follows: 94°C for 3 min, 35 cycles (94°C for 45sec, 57°C/58°C(lamininα5/Itgα6b) for 30sec, 72°C for 1min), 72°C for 7min, 12°C indefinitely. The primers used were as follows:

lama5tc17 (forward) : 5’ – GGCAACACGCCCTGTAAATA – 3’

lama5tc17 (reverse) : 5’ – CTGCACGCTCACTCATTCGG – 3’

Itga6bfu36 (forward) : 5’ – CGGGGCTGTATTAGCAGGTA – 3’

itga6bfu36 (reverse) : 5’ – GTAGTGAGGCGCGACGTTAT – 3’\

with 506bp band for lama5tc17 and 565bp bands for itga6bfu36 post PCR amplification. lama5tc17mutant allele was identified by sequencing/chromatogram analysis of purified PCR products. lama5tc17mutant allele has a single base pair A to T mutation resulting in premature stop codon. itga6bfu36 mutant allele was identified by PCR product digest (MboI restriction enzyme) at 37°C for 2h, followed by agarose gel run. Homozygous wild types show 2 bands (353, 212bp), heterozygotes show 3 bands (565, 353, 212bp), and homozygous mutants show 1 band (565bp).

### Immunostaining

The embryos were fixed in 4% PFA in PBS for 30min at RT followed by 4°C overnight. This was followed by 2 PBS washes (5min each), methanol upgrade (30%, 50%, 70% in PBS) for 10min each on rocker, 100% methanol for 30min, replaced with fresh 100% methanol, incubated at -20° C overnight. Methanol downgrade (70%, 50%, 30% in PBS) for 10min each on rocker was performed, followed by 2 PBS washes (5min each), 5 PBST(0.8% TritonX-100 in PBS) washes for 10min each. The embryos were blocked in 10% NGS [005-000-121, Jackson Immuno Research Labs] in PBST for 3-4hours on rotor at RT, followed by primary antibody incubation in (1% NGS in PBST) solution using following dilutions: 1:100 E-cadherin [610182, BD Bio transductions] 1:400 Lgl2 (ref), 1:500 PKC-ζ(C-20) [sc-216, Santa Cruz], 1:50 anti-BrdU (SM1667P, Acris Antibodies). They were given 5 PBST washes (30min each), incubated in secondary antibody using following dilutions :1:750 goat anti-mouse Cy3/goat anti-rabbit Cy3/goat anti-rat Cy3[115-165-146; Jackson ImmunoResearch/111-165-144, Jackson ImmunoResearch/112-165-167, Jackson ImmunoResearch Labs)], 1:250 anti-rabbit Alexa 488 conjugated [A11034, Invitrogen] for 3hours at RT on rotor. They were given 5 PBST washes (15min each), fixed in 4% PFA in PBS for at least 30min at RT, given 2 PBS washes (5min each) followed by glycerol upgrade (30%, 50%, 70% in PBS) and used for imaging. The embryos were stained for microridges using 1:500 dilution of Rhodamine Phalloidin (R415, Molecular Probes) for 3-4hours after overnight 4% PFA in PBS fixation and 5 PBST washes (10min each). This was followed by 5 PBST washes (15min each), 4%PFA in PBS fixation at RT, 2 PBS washes(5min each), and glycerol upgradation.

### BrdU treatment

42hpf embryos were dechorionated and transferred to 12 well plate with either 10uM BrdU and 2% DMSO in E3 or only 2% DMSO in E3 for 3hours. They were given 3 washes (15min each) followed by 4% PFA in PBS fixation for 30min at RT then 4°C overnight. This was followed by PBS washes, methanol upgrade and downgrade as mentioned above. After PBS washes, they were treated with 4N HCl for 20 min at RT, followed by PBS, PBST washes to process for immunostainings.

### Confocal imaging

All images (unless mentioned otherwise) were acquired from dorsal head epidermis of zebrafish embryos using olympus fv1200 LSM with Plan Apo 60X/NA:1.42 oil immersion objective at 2.4x zoom and 0.28u z-step taken at resolution of 1024x1024. The pinhole was kept 1 Airy Unit. The laser/PMT gain was adjusted for genetic/developmental conditions giving the brightest signal without saturation. The laser/PMT gain was kept constant within each experiment. For BrdU/Lgl2 staining, the zoom was kept 1x. The embryos were screened for GFP fluorescence after mRNA injection of caax-gfp using mercury lamp of epifluorescent microscope. They were sorted and used for confocal live imaging accordingly. For live imaging, embryos of specified genotype were injected using 100ng/uL CAAX GFP mRNA at 1-4 cell stage. They were screened and sorted into GFP expressing embryos at 2dpf, immobilised using 0.8% low melting agarose [Sigma, A9414-100G] (ref) or 1X E3 in tricain (0.04%) followed by imaging. The laser/PMT gain was adjusted to get the brightest signal in the head epidermis (either 2.4x or 1x zoom was used).

### Quantification and statistical analysis

E-cadherin/ Lgl2 levels, apical perimeter, and cell heights were quantified as described previously (Arora et al., 2020). Briefly, 7 pixel width segmented line tool covering the cell membrane in each z-stack was used to quantify mean intensity and apical perimeter in both periderm and basal epidermis. Cell height was measured by counting the no. of z- slices showing E-cadherin/Lgl2 localisation. 5-8 embryos and 5-10 cells per embryo were quantified for each set. At least 3 sets were quantified for each experiment unless mentioned otherwise. Cell counter plugin in Fiji was used to count no. of BrdU positive cells in basal epidermis which was normalised to total no. of cells (marked by Lgl2) in basal epidermis to obtain percentage of proliferating cells. Microridges post F-actin staining was quantified using FIJI/macro script developed by Pinto C. in the lab (Pinto C et al. 2019). Briefly, in Fiji, the images were converted to 8 bit, binary, smooth images; cell boundaries were marked using polygon tool (maximum 5 cells per image), all cells in image were saved in ROI, the ROI was selected and script run on it. The results obtained were used for statistical plotting and comparisons.

Graphpad PRISM10 and Microsoft365 Excel were used for statistical analysis and plotting graphs. An online tool http://shiny.chemgrid.org/boxplotr/ was used to create bean plots. All statistical comparisons are presented in the S14. In the box and bean plot (Fig 2, 3, 6 7), the middle line represents median, upper box and lower box limit represent 1st quartile and third quartile box limits. The upper end and lower end of the vertical line represents 1.5 times the interquartile range or furthest point from the box limit whichever is closest to the box limit, and each point represents data points used to generate box plots. In scatter dot plots(Fig5, 8),the middle line represents the median. In the bean plot (Fig 9), the middle thick line represents the median; density curve which resembles the mirror violin shape, showing where the data points are most concentrated;and rug (or strip plot) show individual data points.

## Supporting information

https://drive.google.com/file/d/1WY0HeEl2QmHfgBLycdU79sS8asHw-3Ga/view?usp=sharing

## Author’s contribution

Experimentation, Data analysis, Phenotype discovery, Optimisation of experiments, Project conceptualisation and interpretation, Manuscript writing.

## Acknowledgements

This work was performed as part of a program of research to understand multi-layered epithelial systems in the Sonawane lab at the Department of Biological Sciences, TIFR Mumbai. Tooba Khan conceptualized and tested the possibility of EMT and tested the role of laminins in this process. The work was conducted using methods and reagents established in the Sonawane lab and was supported by the funds received from TIFR-Department of Atomic Energy (Project Identification no. RTI4003). Tooba Khan was supported by a Ph.D fellowship from TIFR-DAE. The Itga6b (fu36) allele, used in this study, was generated by Dr. Chaitanya Dingare and Prof. Virginie Lecaudey (Institute for Cell Biology and Neuroscience, Goethe University, Germany) and shared with the Sonawane lab.

## Supplementary Information

**Fig. S1.**
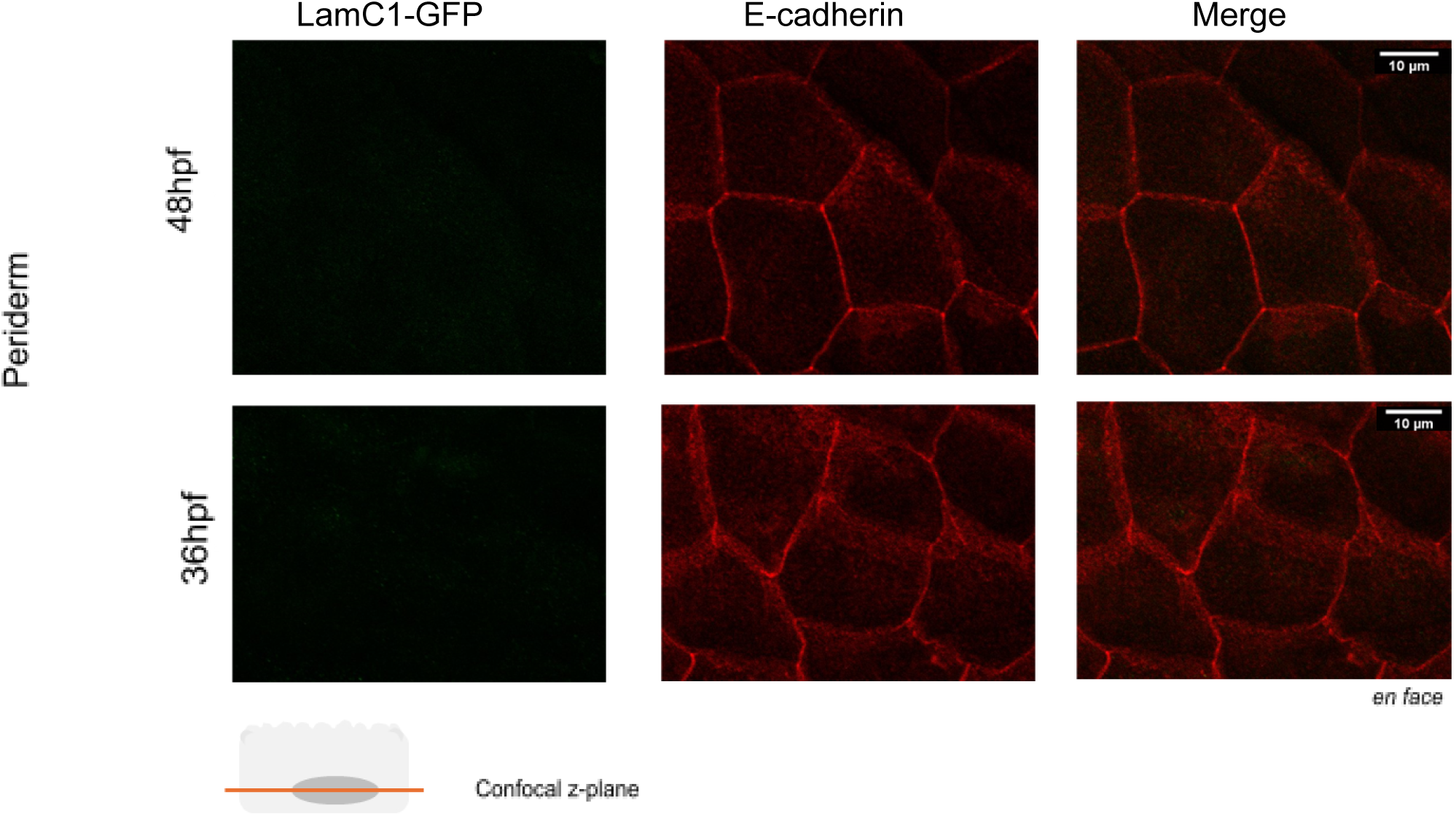
Basal epidermis expresses and is enriched in laminin gamma1 (*lamc1*). LamininC1-GFP was found to be absent in periderm cells marked by E-cadherin, at 48hpf and 36hpf (Fig S1). The schematic diagram depicts one periderm cell. Confocal z-plane lies anywhere inside the cell. (See supplementary Fig. S1). Scale bar depicts 10um length in *en face* images. In images, LamC1-GFP is shown in green LUT, and membrane Lgl2 in red LUT (in Fiji).

Fig S2: Junctional E-cadherin levels and cell morphology is affected in basal epidermis in *lama5* mutants. Source file S2: https://docs.google.com/spreadsheets/d/1fvzG15Fo5xg3eixTTXsDfdGBBBCSl-Ux/edit?usp=sharing&ouid=115433756496797593603&rtpof=true&sd=true

Fig S3: Basolateral membrane polarity protein Lgl2 level is affected in basal epidermis in *lama5* mutants and phenocopies cell morphology effects shown by junctional E-cadherin. Source file S3: https://docs.google.com/spreadsheets/d/13WWs8tUEjKs8-s9jPcHYEB9Dm_wVIZ4z/edit?usp=sharing&ouid=115433756496797593603&rtpof=true&sd=true

Movie S4. Loss of junctional E-cadherin in basal epidermis causes cell-cell detachment with cells throwing dynamic protrusions at cell boundaries. Live movie of CAAX-gfp injected embryos at 1-4 cell stage show intact cell shape and boundaries in *lama5* wt (T=13.44’) while in *lama5* mutants (T=21.12’) CAAX-gfp is diffuse filling up the cells that show cell-cell detachment and throw dynamic protrusions at cell periphery. Duration of live movie for *lama5* wt is 13.44’ (maximum intensity projection of few z-stacks z29-z36, no. of time frames=7, each time frame (T)=3.32’); duration of live movie for *lama5* mut is =21.12’ (single z-stack z44, no. of time frames=9, each time frame (T)=3.32’). WT sib movie S2 (basal epidermis): https://drive.google.com/file/d/1KquUvaIUpkvU2p1yJdfSdKONaNKV0R2h/view?usp=sharing *lama5* movie S2 (basal epidermis): https://drive.google.com/file/d/1mvkd0vRsB9spPka3U7PsOJlUvrIeuNf6/view?usp=sharing

Fig S6: Junctional E-cadherin levels and cell morphology is affected in periderm in *lama5* mutants. Source file S6: https://docs.google.com/spreadsheets/d/1Lm3mH8Q-d7fusMc2rvqqH6oD5QRc20Wb/edit?usp=sharing&ouid=115433756496797593603&rtpof=true&sd=true

Fig S7: Basolateral membrane polarity protein Lgl2 level is not significantly affected in periderm in *lama5* mutants and phenocopies cell morphology effects shown by junctional E-cadherin.Source file S7: https://docs.google.com/spreadsheets/d/1bO9KauoPzAU_Ih3tr6fC9Egveo28u14h/edit?usp=sharing&ouid=115433756496797593603&rtpof=true&sd=true

Fig S9 F-actin based apical polarity structures microridges in periderm form smaller and less complex patterns in *lama5* mutants. Source folder S9: https://drive.google.com/drive/folders/1HoJ30dcmuLSfP8CBiGYUq09MtCtUPKOc?usp=sharing

Movie S10. Periderm shows intact cell-cell adhesion and cell shape in *lama5* mutants. Live movie of CAAX-gfp injected embryos at 1-4 cell stage show intact cell shape and boundaries in *lama5* wt and *lama5* mutants. Duration of live movie for *lama5* wt is 8.53’ (maximum intensity projection of few z-stacks z1-z19, no. of time frames=8, each time frame (T)=3.32’); duration of live movie for *lama5* mut is 12.37’ (maximum intensity projection of few z-stacks z1-z29, no. of time frames=8, each time frame (T)=3.32’). WT sib movie S3 (periderm): https://drive.google.com/file/d/1WMQ73jptjk-LA5c3UyNQaVb75e0aLkYp/view?usp=sharing *lama5* mut movie S3 (periderm): https://drive.google.com/file/d/1OUIP1RY8AV-ULKipQQRu1fAuOSezWGVw/view?usp=sharing

Fig S11: Junctional E-cadherin levels and cell morphology is affected in basal epidermis in *itga6b* and *lama5;itga6b* like *lama5*. Source file S11: https://docs.google.com/spreadsheets/d/1Z0xEbtQvOnn64iCNBHKy_FLI4TCHDXe_/edit?usp=sharing&ouid=115433756496797593603&rtpof=true&sd=true

**Fig. S12.**
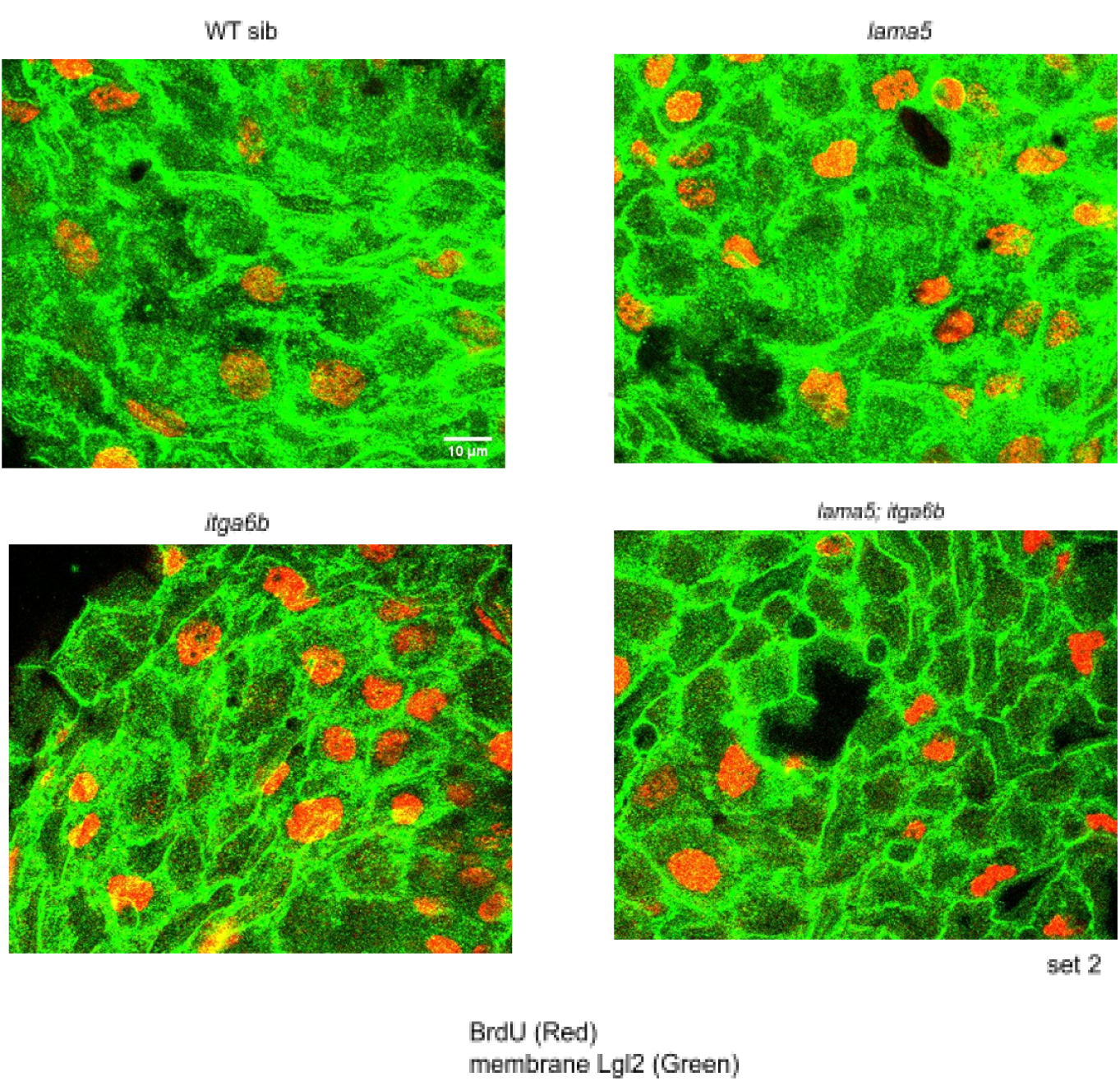
Basal epidermis shows higher proliferation marked by BrdU positive cells in *lama5* and/or *itga6b* mutants. There is an increase in BrdU positive cells in basal epidermis of *lama5* and/or *itga6b* mutants seen across two sets. (Fig 12, Fig S12). Scale bar depicts 10um length. In images, BrdU is shown in red LUT, and membrane Lgl2 in green LUT (in Fiji).

**Fig. S13.**
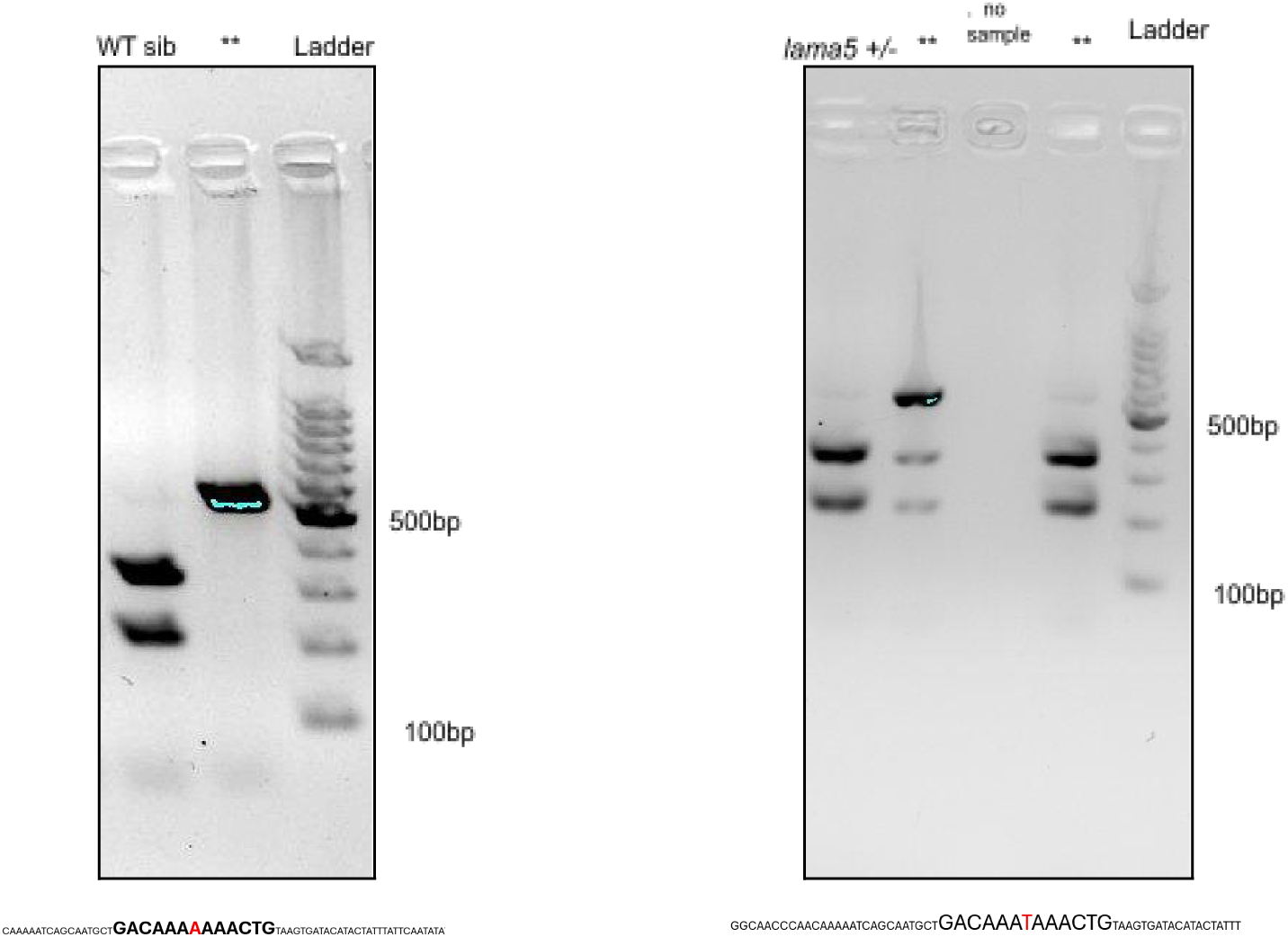
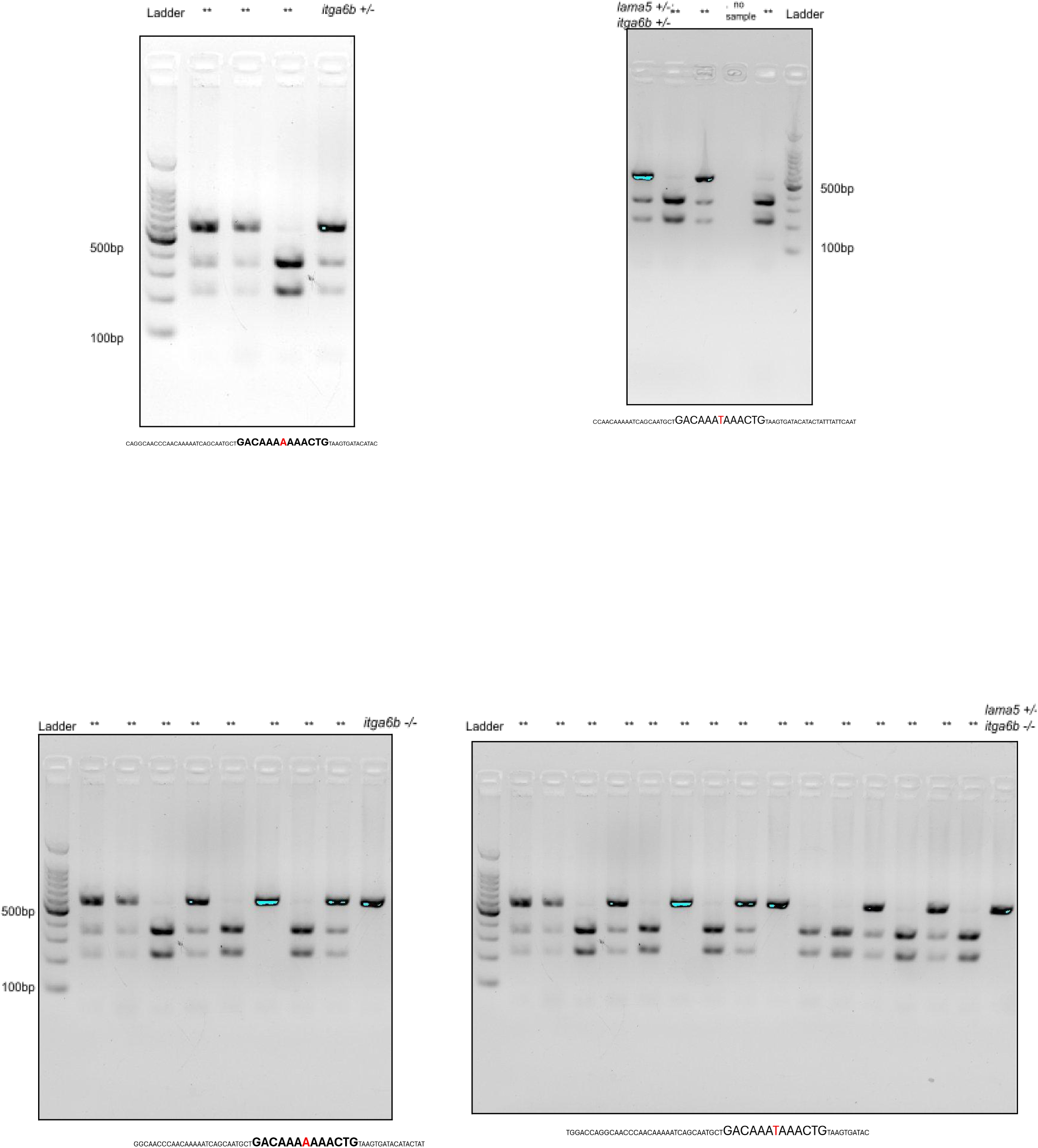
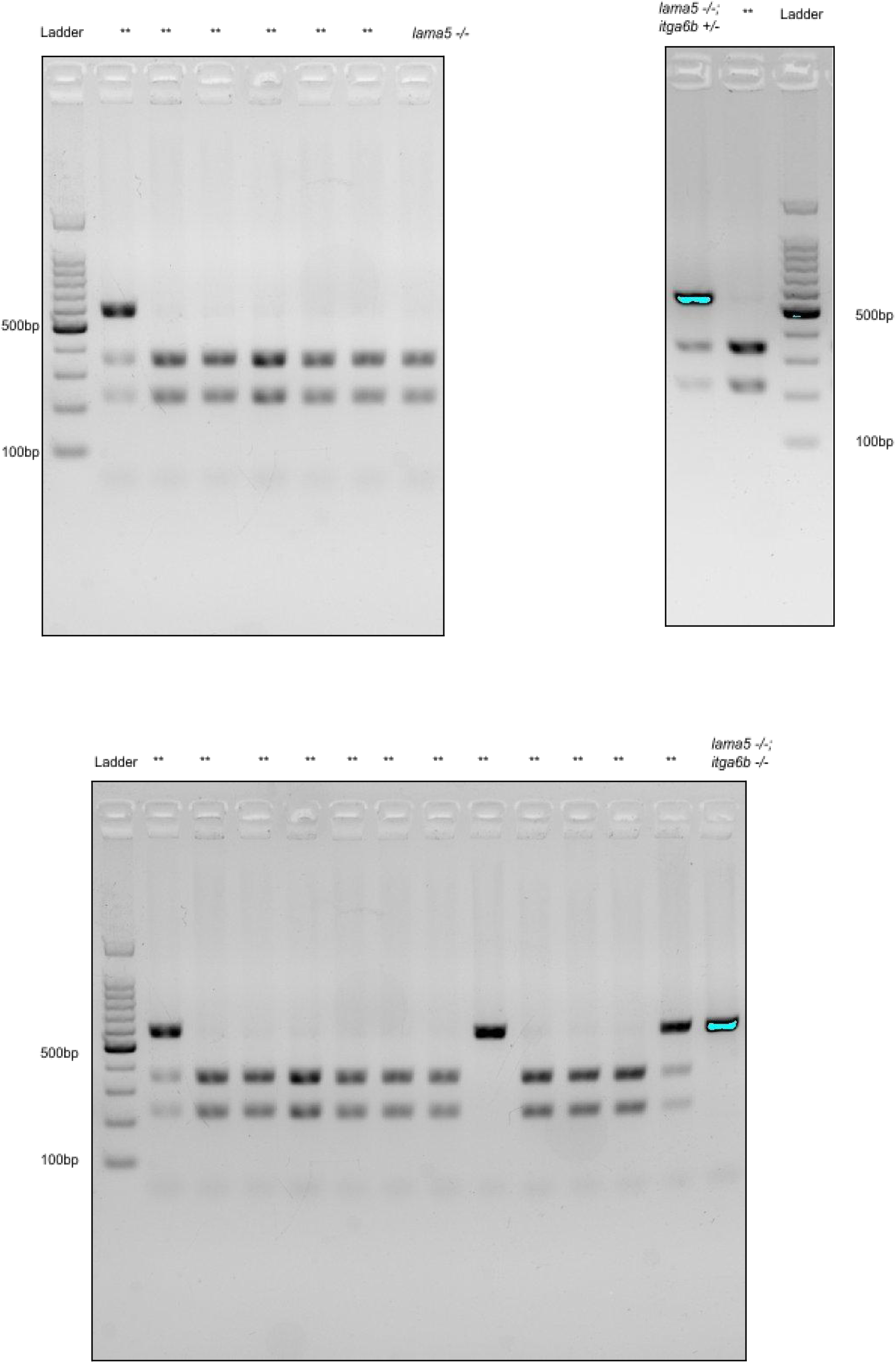
Loss of junctional E-cadherin in basal epidermis causes cell-cell detachment with cells throwing dynamic protrusions at cell boundaries upon single and/or double loss of *lama5* and/or *itga6b* allele. Agarose gel images of PCR product digest (MboI restriction enzyme) are used to identify *itga6b* allele; while sequencing/chromatogram analysis of purified PCR products was used for identification of *lama5^tc17^* heterozygote alleles in WT sib. *itga6b^fu36^* mutant allele- Homozygous wild types show 2 bands (353, 212bp), heterozygotes show 3 bands (565, 353, 212bp), and homozygous mutants show 1 band (565bp). *lama5^tc17^* heterozygotes show a single base pair **<u>A</u>** to **<u>T</u>** mutation resulting in premature stop codon given below the gel images. The double asterisk (**) in gel image denotes ‘other samples; ‘no sample’ denotes empty wells.

Fig S14: Figure wise numerical data & statistical analysis. https://docs.google.com/spreadsheets/d/1s5Qst3eosg53MMAo1deW0Oo0wzFfoToO/edit?usp=sharing&ouid=115433756496797593603&rtpof=true&sd=true

