## Supplementary material for "Unraveling the role of laminin(*lama5*) in maintenance of epithelial identity and polarity in bilayer zebrafish epidermis during development": https://drive.google.com/file/d/1WY0HeEl2QmHfgBLycdU79sS8asHw-3Ga/view?usp=sharing

**Supplementary Information**

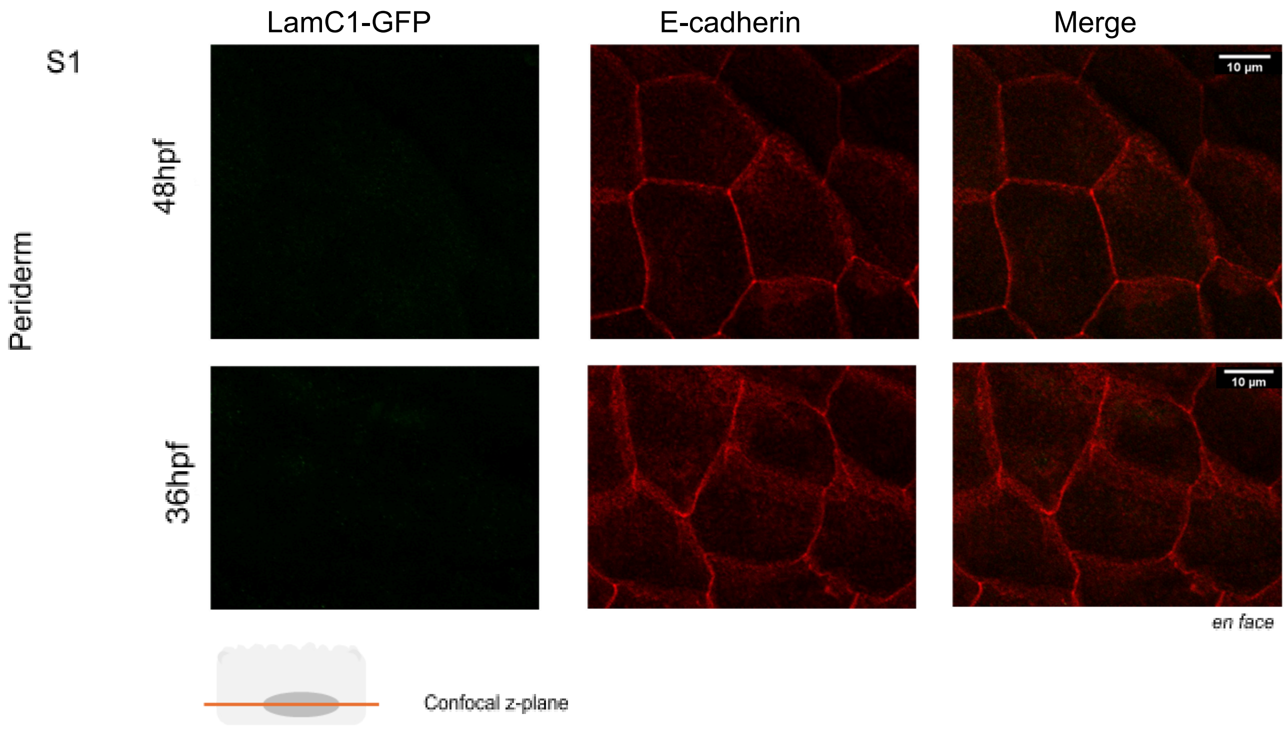


Fig. S1. Basal epidermis expresses and is enriched in laminin gamma1 (*lamc1*). LamininC1-GFP was found to be absent in periderm cells marked by E-cadherin, at 48hpf and 36hpf (Fig S1). The schematic diagram depicts one periderm cell. Confocal z-plane lies anywhere inside the cell. (See supplementary Fig. S1). Scale bar depicts 10um length in *en face* images. In images, LamC1-GFP is shown in green LUT, and membrane Lgl2 in red LUT (in Fiji).

*lama5* mut movie S3 (periderm):
https://drive.google.com/file/d/1OUIP1RY8AV-ULKipQQRu1fAuOSezWGVw/view?usp=sharing

Fig S11: Junctional E-cadherin levels and cell morphology is affected in basal epidermis in *itga6b* and *lama5;itga6b* like *lama5*. Source file S11:
https://docs.google.com/spreadsheets/d/1Z0xEbtQvOnn64iCNBHKy_FLI4TCHDXe_/edit?usp=sharing&ouid=115433756496797593603&rtpof=true&sd=true

S12

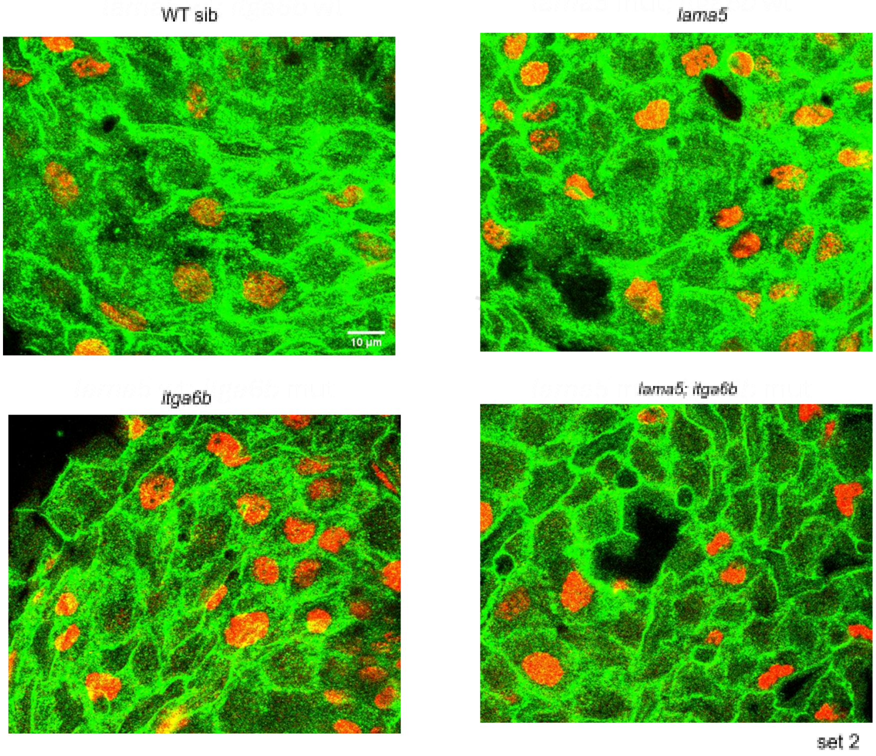


BrdU (Red)
membrane Lgl2 (Green)

Fig. S12. Basal epidermis shows higher proliferation marked by BrdU positive cells in *lama5* and/or *itga6b* mutants. There is an increase in BrdU positive cells in basal epidermis of *lama5* and/or *itga6b* mutants seen across two sets. (Fig 12, Fig S12). Scale bar depicts 10um length. In images, BrdU is shown in red LUT, and membrane Lgl2 in green LUT (in Fiji).

S13

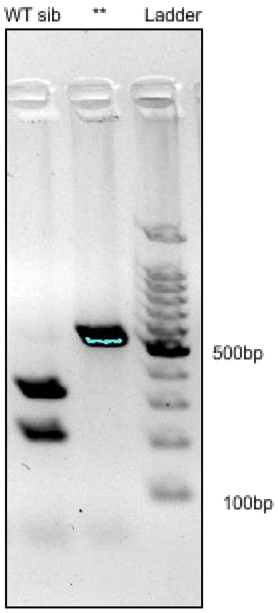


CAAAAATCAGCAATGCT**GACAAAAAAACTG**TAAGTGATACATACTATTTATTCAATATA

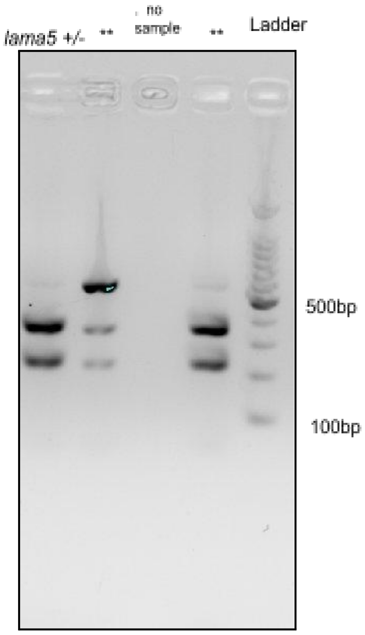


GGCAACCCAACAAAAATCAGCAATGCTGACAAATAAACTGTAAGTGATACATACTATTT

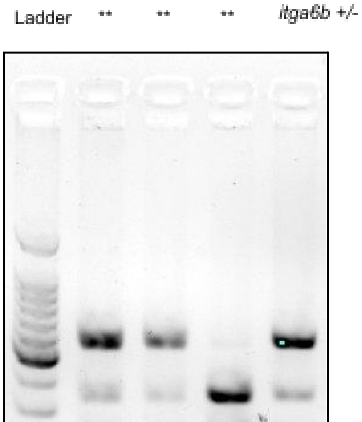


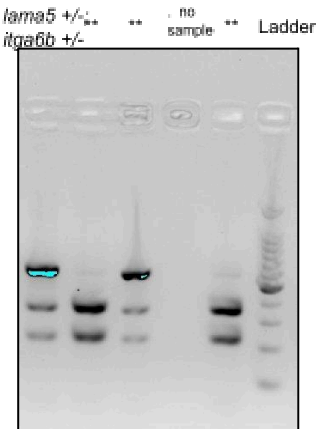

500bp

100bp

CAGGCAACCCAACAAAAATCAGCAATGCT**GACAAAAAAACTG**TAAGTGATACATAC

500bp

100bp

CCAACAAAAATCAGCAATGCTGACAAATAAACTGTAAGTGATACATACTATTTATTCAAT

Ladder ** ** ** ** ** ** ** ** *itga6b -/-*

500bp

100bp

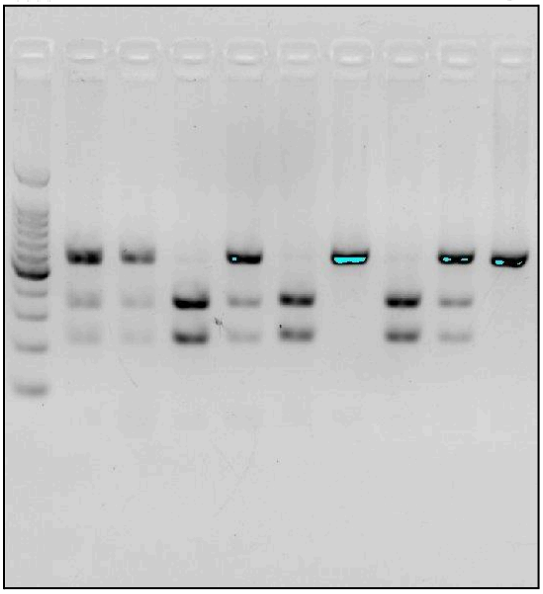

GGCAACCCAACAAAAATCAGCAATGCT**GACAAAAAAACTG**TAAGTGATACATACTAT

Ladder ** ** ** ** ** ** ** ** ** ** ** ** ** ** ** *lama5 +/-* *itga6b -/-*

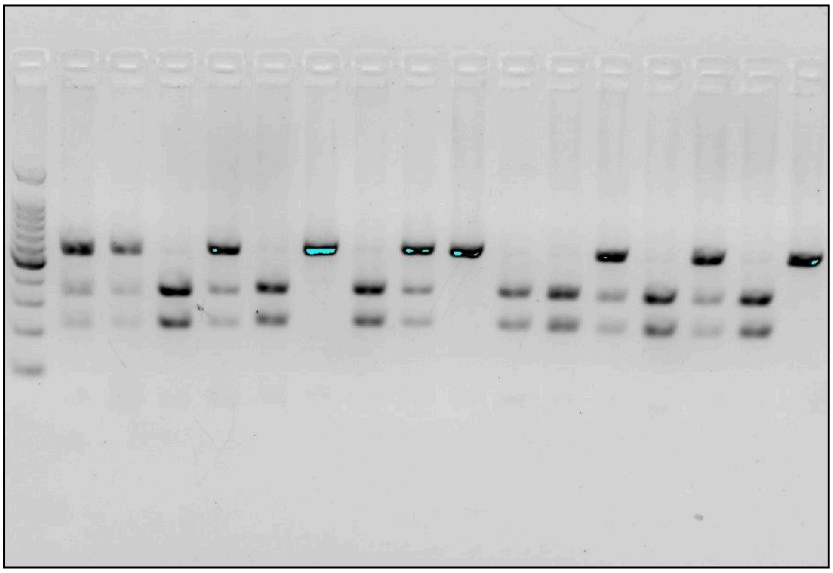

TGGACCAGGCAACCCAACAAAAATCAGCAATGCTGACAAATAAACTGTAAGTGATAC

Ladder ** ** ** ** ** ** *lama5 -/-*

500bp

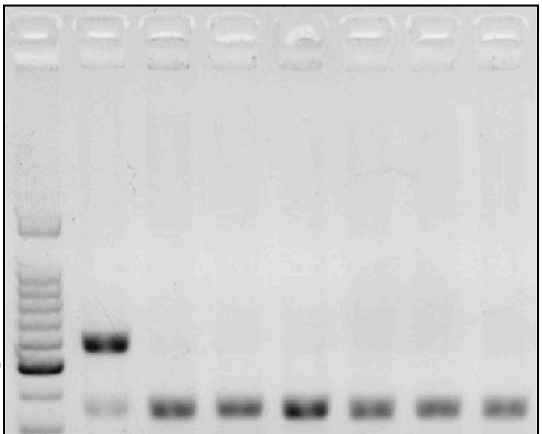

*lama5 -/-;* *itga6b +/-* ** Ladder

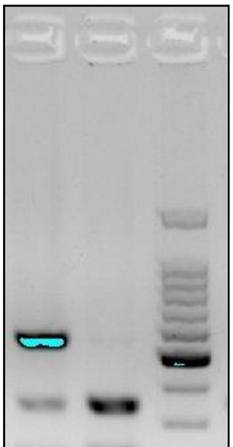

500bp

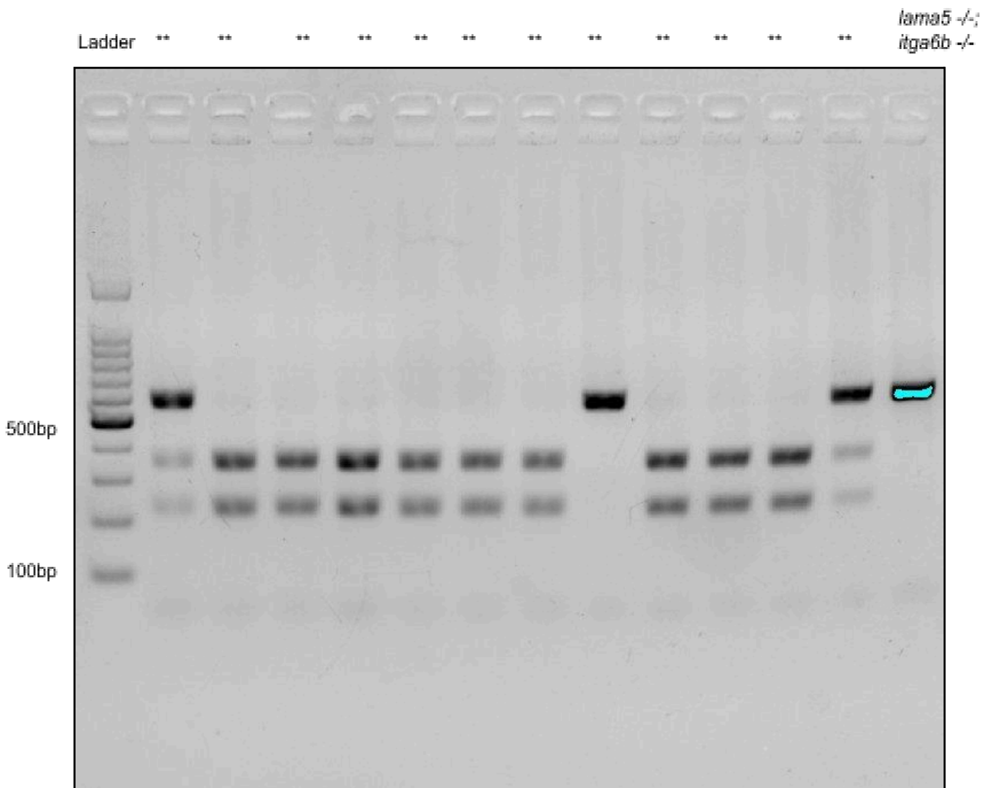
Ladder
**
**
**
**
**
**
**
**
**
**
**
**
lama5 -/-;
itga6b -/-
500bp
100bp

Fig. S13. Loss of junctional E-cadherin in basal epidermis causes cell-cell detachment with cells throwing dynamic protrusions at cell boundaries upon single and/or double loss of *lama5* and/or *itga6b* allele. Agarose gel images of PCR product digest (MboI restriction enzyme) are used to identify *itga6b* allele; while sequencing/chromatogram analysis of purified PCR products was used for identification of *lama5*$^{tc17}$ heterozygote alleles in WT sib. *itga6b*$^{fu36}$ mutant allele- Homozygous wild types show 2 bands (353, 212bp), heterozygotes show 3 bands (565, 353, 212bp), and homozygous mutants show 1 band (565bp). *lama5*$^{tc17}$ heterozygotes show a single base pair **A** to **T** mutation resulting in premature stop codon given below the gel images. The double asterisk (**) in gel image denotes 'other samples; 'no sample' denotes empty wells.

Fig S14: Figure wise numerical data & statistical analysis.
https://docs.google.com/spreadsheets/d/1s5Qst3eosg53MMAo1deW0Oo0wzFfoToO/edit?usp=sharing&ouid=115433756496797593603&rtpof=true&sd=true
